# Long-term T cell fitness and proliferation is driven by AMPK-dependent regulation of oxygen reactive species

**DOI:** 10.1101/2020.03.06.978791

**Authors:** Anouk Lepez, Tiphène Pirnay, Sébastien Denanglaire, David Perez-Morga, Marjorie Vermeersch, Oberdan Leo, Fabienne Andris

**Affiliations:** Laboratory of Immunobiology, Université Libre de Bruxelles, Belgium; Center for Microscopy and Molecular Imaging (CMMI), Université Libre de Bruxelles, Belgium; Laboratory of Molecular Parasitology, Université Libre de Bruxelles, Belgium

**Keywords:** T lymphocytes, cell metabolism, homeostasis, ROS

## Abstract

The AMP-activated kinase (AMPK) is a major energy sensor metabolic enzyme that is activated early during T cell immune responses but its role in the generation of effector T cells is still controversial.

Using both *in vitro* and *in vivo* models of T cell proliferation, we show herein that AMPK is dispensable for early TCR signaling and short-term proliferation but required for sustained long-term T cell proliferation and effector / memory T cell survival. In particular, AMPK promoted accumulation of effector / memory T cells in competitive homeostatic proliferation settings. Transplantation of AMPK-deficient hematopoïetic cells into allogeneic host recipients led to a reduced graft-versus-host disease, further bolstering a role for AMPK in the expansion and pathogenicity of effector T cells.

Mechanistically, AMPK expression enhances the mitochondrial membrane potential of T cells, limits reactive oxygen species (ROS) production, and resolves ROS-mediated toxicity. Moreover, dampening ROS production alleviates the proliferative defect of AMPK-deficient T cells, therefore indicating a role for an AMPK-mediated ROS control of T cell fitness.

## Introduction

Changes in the proliferative and functional state of T lymphocytes during an immune response are accompanied by profound modifications in the cell metabolism. Whereas naive T cells generate the largest part of their ATP through mitochondrial oxidative phosphorylation (OxPhos), activated lymphocytes rely heavily on glycolysis to fulfill their energetic demands (1, 2). Long-lived memory T cells that subsist after primary infection convert back to an oxidative metabolism with a marked preference for fatty acid as primary energy source (FAO) (1, 3).

The AMP-activated protein kinase (AMPK) maintains the energy balance in eukaryotic cells and is a central regulator of cellular responses to metabolic stress (4–6). Activation of AMPK decreases fatty acid and protein synthesis (7, 8), and promotes FAO and mitochondrial biogenesis (9, 10). AMPK also up-regulates the rate of glycolysis through phosphorylation of PFKFB3 and increases glucose uptake (11, 12). Moreover, AMPK recently emerged as an important sensor and regulator of mitochondrial ROS and a regulator of ROS neutralization by antioxidant cellular systems, thereby maintaining adequate cellular oxidative balance (13).

Although T lymphocyte activation results in the rapid activation of AMPK *in vitro* and *in vivo* (3, 14), the role of this enzyme in T cell function is still controversial. AMPK appears dispensable for the generation of effector T cells in two independent studies (15, 16) while required for the expansion of antigen-specific CD8^+^ T cells in response to viral or bacterial infections (17). Moreover, effector T cells lacking AMPKα1 display reduced mitochondrial bioenergetics and cellular ATP in response to glucose limitation, in support of a role for AMPK in the metabolic adaptations of activated effector T cells (17). Despite these observations, mice lacking expression of AMPK in the T cell compartment appear as largely immunocompetent, thus questioning the specific role of this enzyme in regulating immune homeostatis.

We confirm in this report that AMPK is dispensable during the earliest steps of T cell activation. AMPK-deficient cells however display reduced capacity to sustain long term proliferative responses both *in vitro* and *in vivo* revealing a potentially important role for this enzyme in the setting of continuous stimulation. We further demonstrate that AMPK expression protects cells from the damaging effects of reactive oxygen species (ROS) that accumulate during rapid cell division, thus revaling the important role of this metabolic sensor in ensuring long term proliferative T cell fitness.

## Results

### AMPK promotes effector / memory T cells engraftment in the peripheral tissues of bone marrow chimeras

To investigate the role of AMPK in T cell homeostasis, we used conditional KO mice in which loxP-flanked alleles of AMPKα1 (prkaa1^f/f^) are deleted by the Cre recombinase expressed from the T-cell-specific cd4 promoter (prkaa1^f/f^ - CD4-Cre mice, hereafter referred as AMPK^KO-T^ mice). No major abnormalities in the thymic and splenic lymphoid populations of these mice have been reported (15, 16). To evaluate whether expression of AMPK modulates the size of T cell compartments in a homeostatic competitive environment, we generated chimeras by reconstituting irradiated CD3ε^−/−^ recipient mice with a 1:1 mixture of congenitally marked bone marrow cells from wild type (CD45.1^+^) and AMPK^KO-T^ (CD45.2^+^) donor mice. The average contribution of WT and AMPK-KO cells to the repopulation of the thymus, spleen and lymph nodes CD4 and CD8 T cell compartments was comparable (Figure 1A, B). Of interest, AMPK-KO splenocytes expressed slightly but significantly lower percentages of CD44^hi^ CD62L^low^ surface markers characteristic of effector / memory cells (Figure 1C-F). The AMPK-KO T cell compartment was also markedly deficient in the expression of the tissue homing CXCR3 chemokine receptor (Figure 1G, H). Accordingly, T cells derived from AMPK^KO-T^ mice contributed only marginally to the repopulation of the gut lymphoid tissues when compared to wild type T cells three months after competitive bone marrow transplantation (Figure 1A, B and supplemental Figure S1A). A reduced reconstitution of the AMPK-KO T cell pool was also observed in the liver of chimeras (supplemental Figure S1B). Thus, AMPK is required for T cells to properly exit from naive “progenitor” state and could provide a competitive advantage to effector / memory phenotype T cells in engrafting the niche of non-hematopoietic tissues following adoptive transfer of bone marrow precursors in T cell deficient mice.

**Figure 1:**
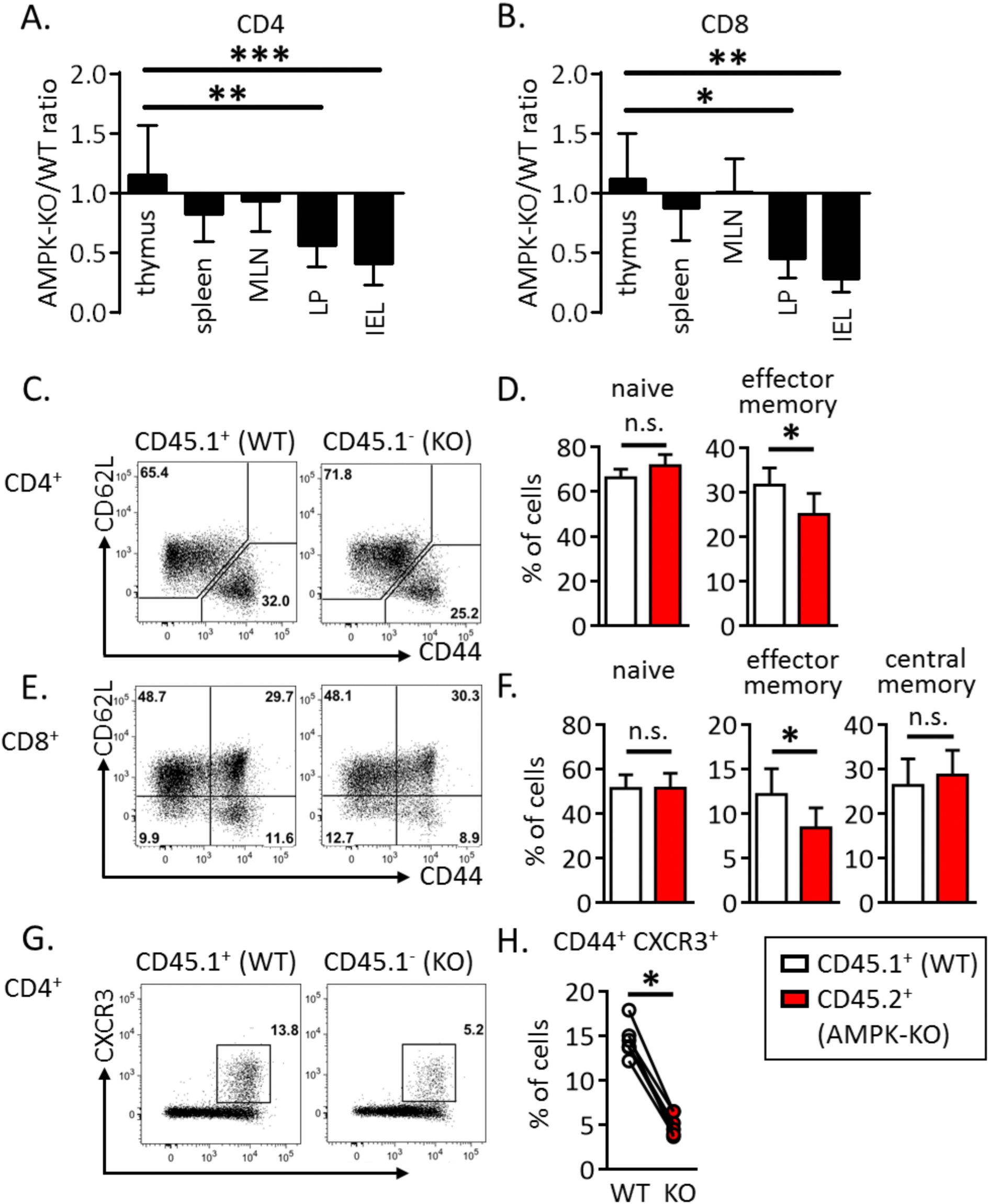
AMPK promotes peripheral T cells replenishment upon bone marrow transfer. (**A-B**) Relative contribution of AMPK-KO CD4^+^ (**A**) and CD8^+^ (**B**) T cells in the thymus, spleen, MLN, LP and IEL of CD3ε^-/-^ mice were reconstituted with a 1:1 mixture of bone marrow cells from WT (CD45.1) and AMPK^KO-T^ (CD45.2) mice. Data are expressed as a ratio of AMPK-KO over WT cells in the CD4^+^ or CD8^+^ T cell gate. (**C-H**) CD62L, CD44 and CXCR3 expression by WT and AMPK-KO T cells recovered from the spleen of mice reconstituted as in (**A, B**). Representative dot plots (**C, E**) and histograms showing the mean % +/- SD (**D, F**) in different cell gates as indicated. Data are representative of 3 (**A, C, D, G, H**) or 2 (**B, E, F**) independent experiments with n = 6. Statistical analysis: Friedman followed by Dunn multiple comparison (**A, B**), Mann–Whitney (**D, F**), Wilcoxon Test (**H**). * p <0.05 ; ** p < 0.01 ; *** p < 0.001

### AMPK promotes T cell homeostatic proliferation and confers higher pathogenicity to donor T cells during GVHD

We next used a well-established model of homeostatic proliferation to further study the long-term / sustained T cell proliferation of AMPK-KO CD4^+^ T cells (18). CFSE-labeled, congenitally marked, CD44^low^ CD4^+^ T cells were purified from WT and AMPK^KO-T^ mice and co-transferred into CD3ε^−/−^ lymphopenic recipient mice. Ten days after adoptive transfer, splenic CD4^+^ T cells expressing high and low CFSE contents, respectively indicative of slow and fast homeostatic proliferation, were recovered in both WT and AMPK-KO subsets. As expected, fast divided cells up-regulated CD44 expression (supplemental Figure S2A). Despite starting with similar number of cells, the recovery of donor WT CD4^+^ T cells was reproducibly greater than that of AMPK-KO cells in each organ tested (Figure 2A, B). Co-transfer of naive B6.CD45.1 and B6.CD45.2 (both WT genotype) led to similar CD4^+^ T cell recovery, indicating that CD45 polymorphism does not affect homeostatic proliferation of T lymphocytes (supplemental Figure S3A, B). Of interest, subdivision of splenic CD4^+^ T cell populations based on CFSE content indicated that the relative contribution of AMPK-KO cells to the T cell pool decreased over T cell divisions (Figure 2C, D; see also supplemental Figure S2B for individual mouse representation). The lower percentage of AMPK-KO T cells undergoing fast division rates was further reflected by their lower expression of the Ki67 proliferation marker (Figure 2E). Wild type fast dividing cells acquired potential for IFNγ secretion as well as cell surface expression of CXCR3 (Figure 2F, H, upper panels: compare 0-2 and 3 division groups). Both CXCR3 expression and IFNγ secretion were significantly reduced in the AMPK-KO subset whereas comparable IL-2 secretion was observed in WT and KO groups (Figure 2F-J).

**Figure 2:**
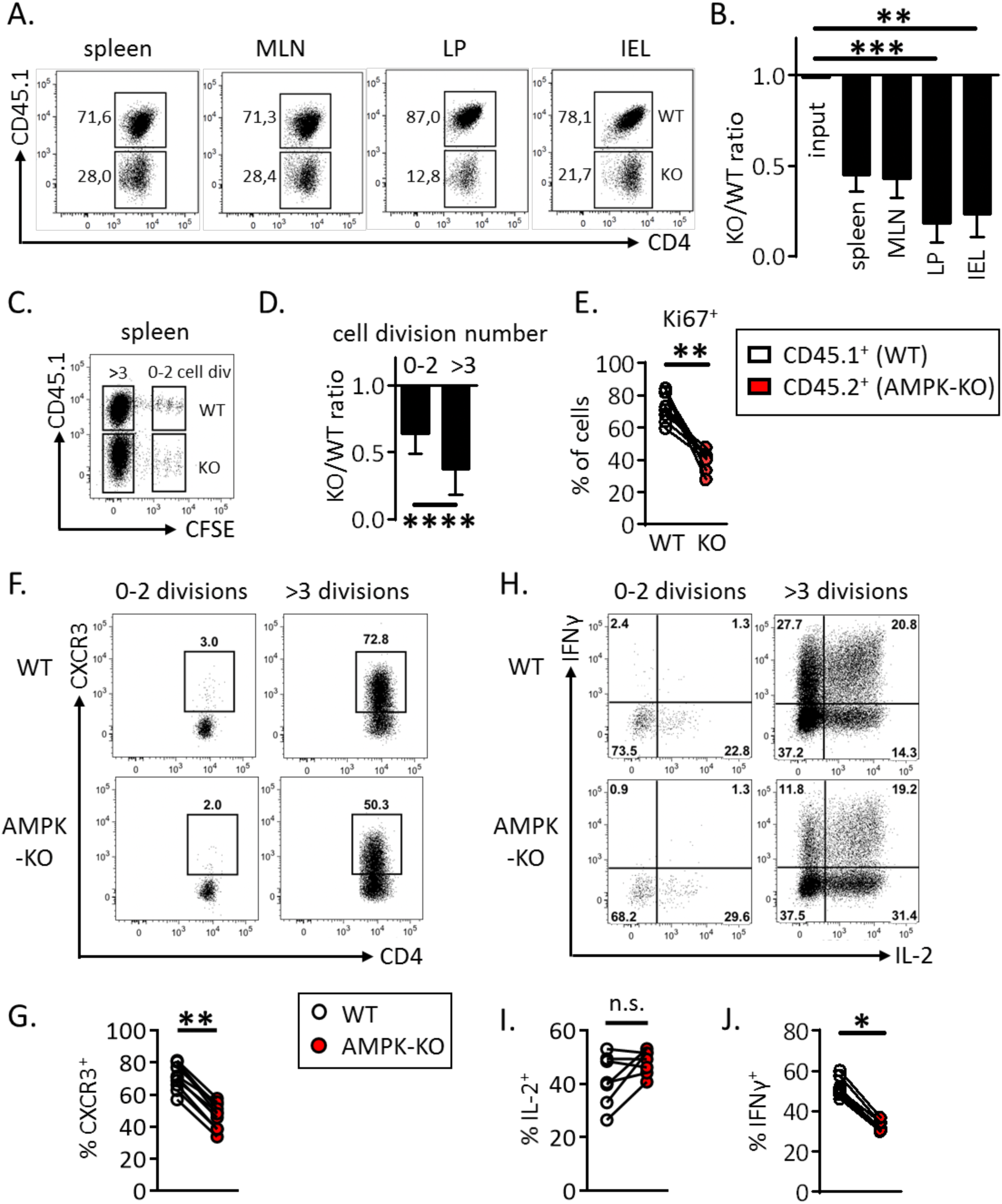
WT CD4^+^ T cells outcompete their AMPK-KO counterparts during homeostatic proliferation. CFSE labeled CD4^+^ naive T cells from WT (CD45.1) and AMPK^KO-T^ (CD45.2) mice (1:1 ratio) were i.v. injected into CD3ε^-/-^ mice. Recovered T cells were analyzed on day 10 by flow cytometry. (**A, B**) Relative contribution of AMPK-KO T cells to the repopulation of the CD4^+^ T cell subset in the spleen, MLN, LP and IEL (CD4^+^ gated). Representative dot plots (**A**) and histograms showing the ratio of AMPK-KO / WT cells (**B**, gate CD4^+^ T cells). Data are representative of 3 (LP, IEL), 4 (MLN) or 6 (spleen) independent experiments with n=6. (**C**) CFSE dilution versus CD45.1 expression by CD4^+^ T cells in the spleen. (**D**) Relative contribution of AMPK-KO T cells in the pool of recovered splenic CD4^+^ T cells that divided 0-2 or > 3 times, according to CFSE dilution (pool of 4 independent experiments with n = 15). (**E**) Percentage of WT and AMPK-KO CD4^+^ T cells that express Ki67 in the spleen (1 experiment representative of 2, with n = 5). (**F-H**) Percentage of WT and AMPK-KO CD4^+^ T cells expressing CXCR3 (**F**) or producing IL-2 and IFNγ (**H**) according to the number of cell divisions in the spleen. (**G-J**) Percentage of WT and AMPK-KO CD4^+^ T cells expressing CXCR3 (**G**), IL-2 (**I**) or IFNγ (**J**) among CD4^+^ T cells that divided more than 3 times (pool of 2-3 independent experiments, with n = 7-10). Statistical analysis: Friedman followed by Dunn multiple comparison (**B**), Wilcoxon (**D, E, G, I** and **J**). * p <0.05 ; ** p < 0.01 ; *** p < 0.001, **** p < 0.0001

Similar results were obtained when *in vitro*-derived memory-like Th1 cells were subjected to homeostatic proliferation (see supplemental Figure S4 for details). Moreover undivided WT and AMPK-KO Th1 cells were recovered at similar frequencies upon transfer into lymphoreplete C57BL6 mice, suggesting that AMPK-KO T cells did not experience increased cell death *in vivo* (supplemental Figure S4B).

To further assess if AMPK promotes *in vivo* proliferation in a non-competitive environment, we used a model of graft-versus host disease (GVHD), in which donor T cells respond to widely distributed alloantigens, leading to T cell mediated pathology (19). To this purpose, we transferred wild type or AMPK-KO T cells (H-2^b^), together with wild type T-depleted bone marrow precursors, into lethally irradiated Balb/c (H-2^d^) recipient mice. Decreased percentages of allogeneic CD4^+^ were recovered from the spleen of recipient mice transplanted with AMPK-KO T cells (Figure 3A). Although the proportion of allogeneic Tregs was very low in this experimental setting, we observed a significant increase in their frequencies in AMPK-KO recipients (Figure 3B). No difference was observed in allogeneic CD8^+^ T cells frequency (Figure 3C). Futhermore GVHD clinical score was decreased in recipients of AMPK-KO T cells (Figure 3D). Accordingly, WT T cells caused rapid lethality in almost all the recipients, whereas AMPK-KO T cells only caused lethality in a fraction of the recipients (Figure 3E), thus suggesting that AMPK activation is required to induce T cell pathogenicity and GVHD.

**Figure 3:**
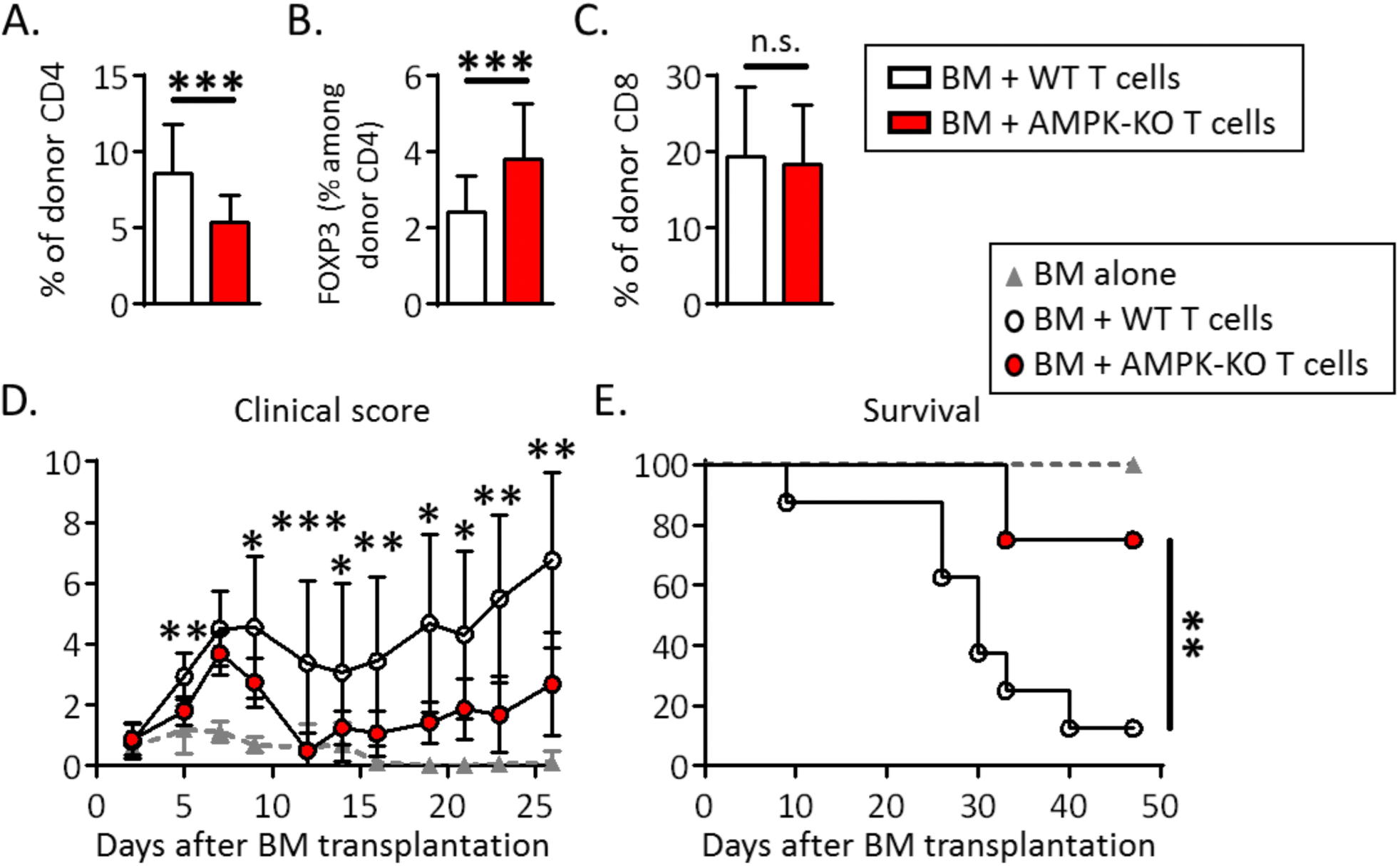
Knock-down of AMPK in donor T cells decreases the severity of GVHD. (**A**-**C**) Percentage of allogenic CD4^+^ (**A**) Tregs (**B**) and CD8^+^ (**C**) T cells in the spleen of the Balb/c host mice. Data are pooled from 5 independent experiments with a total n = 26 (WT group) and 27 (AMPK-KO group). Clinical score (**D**) and survival (**E**) of host mice (n = 8 per group, 1 experiment representative of 3). Statistical analysis: Mann–Whitney (**A-D**), Mantel-Cox (**E**). * p <0.05 ; ** p < 0.01 ; *** p < 0.001

### AMPK is dispensable for IL-7 and TCR early signaling but required for optimal T cell expansion *in vitro*

To further study the role of AMPK in the continuous expansion / survival of CD4^+^ T cells, we developed an *in vitro* model of homeostatic T cell proliferation in the presence of autologous dendritic cells and IL-7, based on a recent report of Rosado-Sánchez et al (20). Two weeks *in vitro* culture of CFSE-labeled naive Th cells with autologous DCs and IL-7, but not DCs or IL-7 alone, led to slow but significant T cell expansion, characterized by increased cell number recovery and the presence of both slow and fast divided cells (Figure 4A and Supplemental Figure S5 for details). As previously observed *in vivo*, only fast dividing cells acquired CD44^hi^ / CD62L^low^ effector cell marker expression and secrete IFNγ (Supplemental Figure S5C). Although a 14-day culture in IL-7-containing media already led to a small decrease in AMPK-KO Th cell recovery, the IL-7+DC-driven proliferation resulted in a further gradual decrease over time (Figure 4B, C). Accordingly, the KO/WT T cell ratio was further decreased among highly divided (CFSE^low^) cells (Figure 4D, E). Similar results have been observed with memory-like Th1 subjected to homeostatic proliferation *in vitro* (Supplemental Figure S6). As previously shown in the *in vivo* model, AMPK-KO Th cells secreted less IFNγ, compared to their wild type counterparts (Figure 4F).

**Figure 4:**
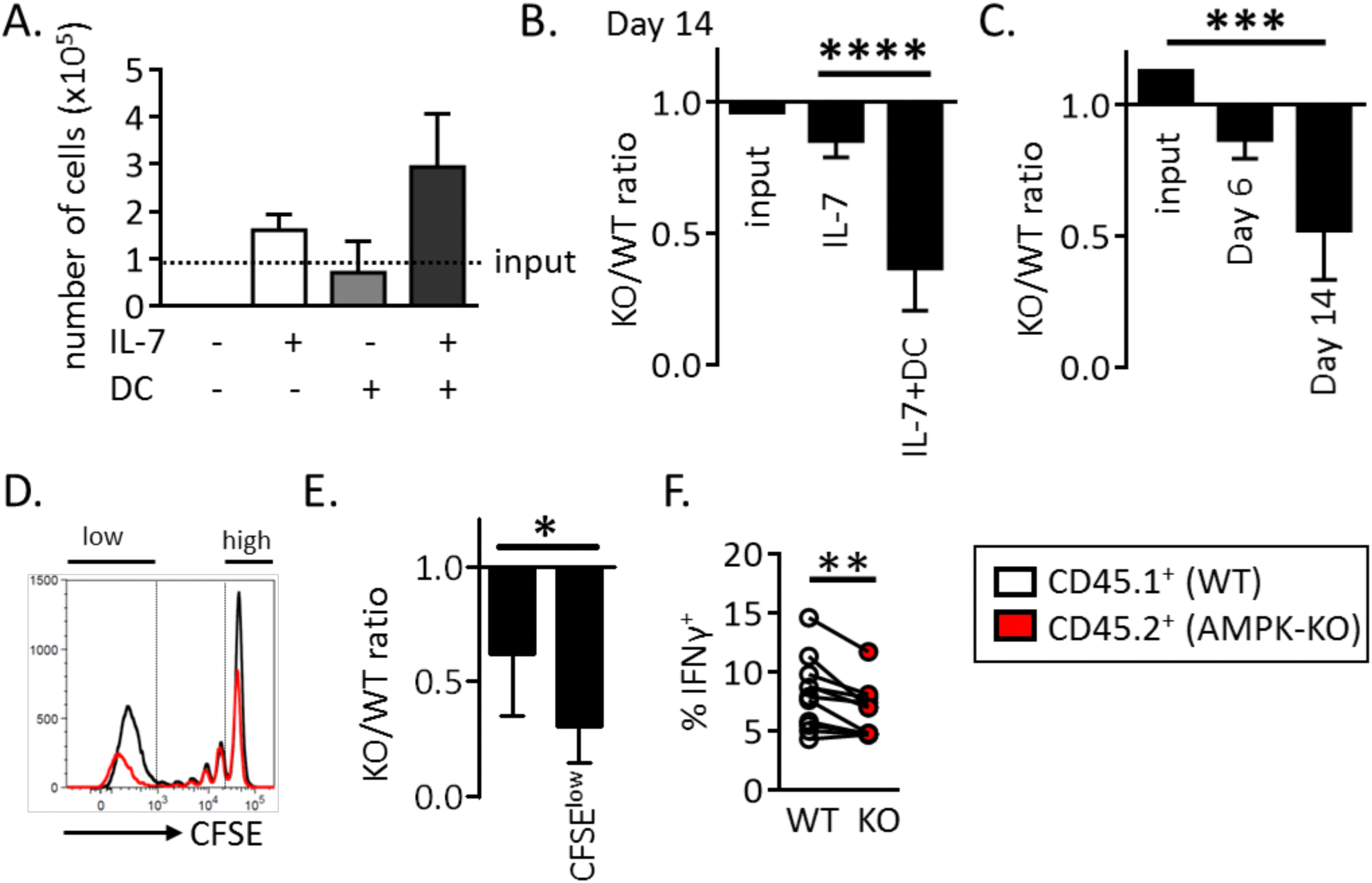
AMPK promotes homeostatic proliferation *in vitro*. (**A**) Naive Th cells from C57BL/6 mice were cultured in the presence of IL-7 and / or syngeneic DCs, as indicated. Culture media were replaced with fresh DCs and IL-7 each 4-5 days. Results indicate the numbers of T cells recovered on day 18, with dashed line corresponding to the number of cells in the input culture (day 0) (data are pooled from 2 independent experiments, n = 3-6). (**B**) Relative recovery of AMPK-KO T cells after 14 day-culture with IL-7 in the presence or absence of syngenic DC. (**C**) Relative recovery of AMPK-KO T cells at day 6 and 14 of IL-7+DC culture. (**D**) CFSE dilution profiles of WT and AMPK-KO at day 14 of IL-7+DC culture. (**E**) Relative contribution of AMPK-KO T cells according to CFSE content. (**F**) Percentage of WT and AMPK-KO T cells expressing IFNγ at day 14. Data are representative of at least 5 experiments with n = 17 (**B**) or 8 (**C**), or are pool of 2 independent experiments with n = 3-6 (**A**), 13 (**E**) or 11 (**F**). Statistical analysis: Mann–Whitney (**B**), Friedman followed by Dunn multiple comparison (**C**) Wilcoxon (**E-F**). * p <0.05 ; ** p < 0.01; *** p < 0.001; **** p < 0.0001

WT and AMPK-KO naive T cells expressed similar levels of phospho-STAT5 and phospho-STAT1 in response to IL-7 and survived, for at least 14 days, in IL-7-supplemented media (supplemental Figure S7 and data not shown), suggesting that defective proliferation does not result from altered IL-7 signaling. In agreement with our previous report (15), AMPK-KO T cells expressed early T cell activation markers and displayed normal short-term TCR-dependent proliferative response to anti-CD3 or allogeneic APCs (supplemental Figure S8A-C). However, co-culture of WT and AMPK-KO T cell in the presence of anti-CD3/CD28 mAbs revealed a small decrease in the AMPK-KO versus WT Th cell recovery. Futhermore WT Th cells largely outcompete their AMPK-KO counterparts when further expanded with IL-2 (supplemental Figure S8D). Collectively, these data indicate that AMPK sustains prolonged cytokine-driven T cell proliferation and/or viability.

### AMPK regulates mitochondrial fitness in T cells

RNA-seq analysis revealed no difference in the pattern of gene expression between WT and AMPK-KO Th cells recovered at the end of a 14-day culture with DCs and IL-7 (Supplemental Figure S9), indicating a negligible role of AMPK in the regulation of gene transcription in this experimental setting.

WT and AMPK-KO CD4^+^ T cells were assessed for mitochondrial bioenergetics. Freshly purified (or rested in IL-7-supplemented media) AMPK-KO naive Th cell expressed a slight reduced mitochondrial membrane potential (Figure 5A and data not shown).This defect was exacerbated in IL-7+DC-expanded T cells and in *in vitro*-derived memory like Th1 (Figure 5B, C and Supplemental Figure S10A, B). Although AMPK-KO T cells recovered after 14 days of homeostatic proliferation exhibited decreased mitochondrial membrane potential, this was not associated with changes in oxygen consumption rate (OCR) or extra-cellular acidification rate (ECAR) (Figure 5D-F). Transmission electron microscopy (TEM) revealed that AMPK-KO T cells did not show difference in mitochondria size and number (Figure 5G-I). Yet, we noticed an increased percentage of mitochondria harboring localized increased membrane density and membrane budding in AMPK-KO cells (Figure 5J, K).

**Figure 5:**
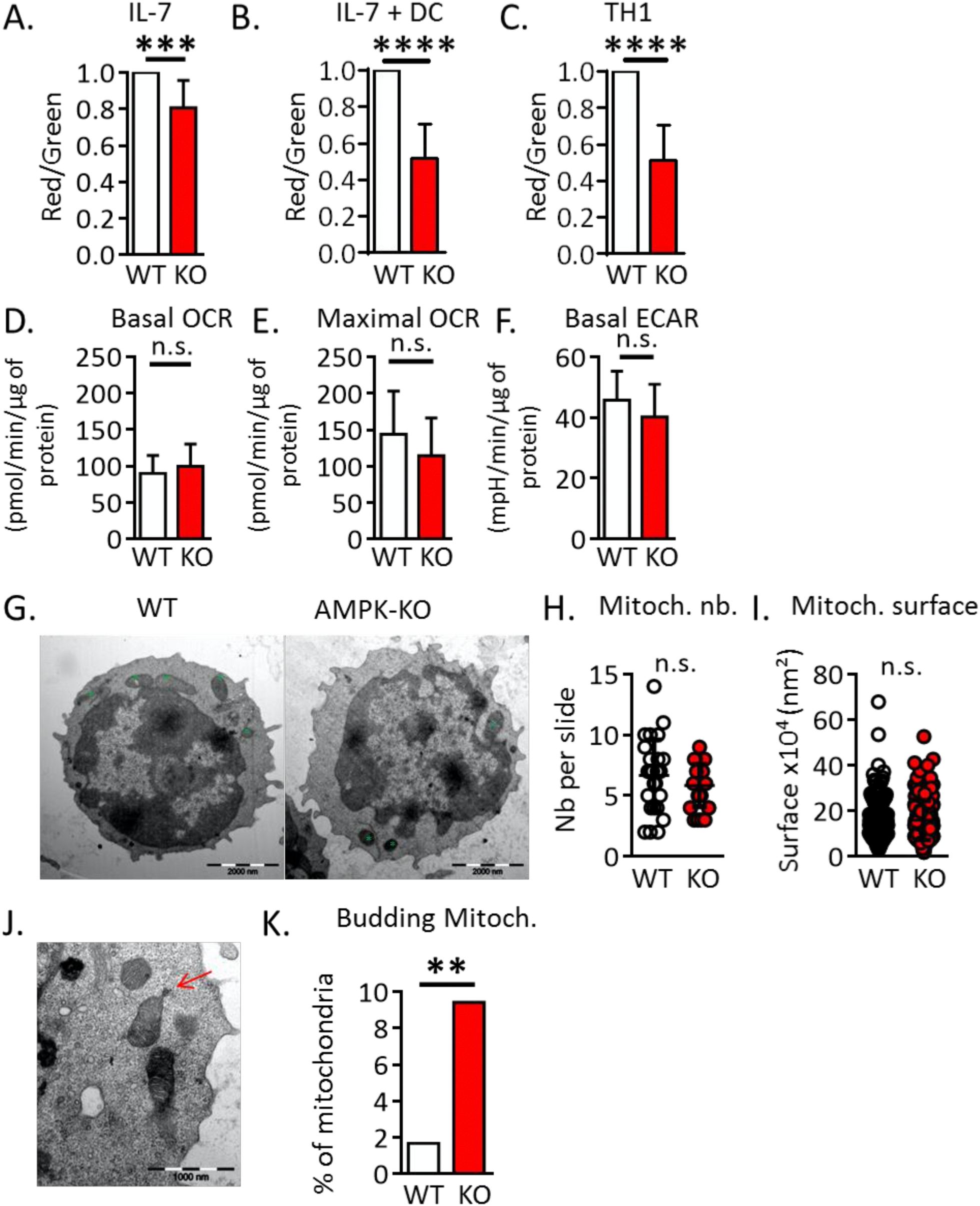
AMPK promotes mitochondrial fitness of T cells. (**A-C**) Mitochondrial membrane potential measured as a ratio of JC1 Red/Green fluorescence ratio in naive T cells (**A**), homeostatic proliferation-recovered T cells (B) and memory-like Th1 cells (**C**), with WT ratio set to 1. (**D**-**F**) Basal OCR (**D**), maximal OCR (**E**) and ECAR (**F**) of WT and AMPK-KO T cells recovered after 14 days of homeostatic proliferation. (**G-I**) TEM analysis of WT and AMPK-KO naive T cells. Representative images (**G**), numbers (**H**) and areas (**I**) of mitochondria (a total of 179 (WT) and 170 (KO) mitochondria from 27 (WT) and 29 (AMPK-KO) T cells analyzed are represented). (**J, K**) Image representative (**J**) and percentage (K) of mitochondria presenting a budding protrusion (WT : 3 out of 179, AMPK-KO : 16 out 170 mitochondria). Data are representative of 7 (**A, B**) or 2 (**H, I, K**) experiments or are pooled from 9 (**C**) or 2 (**D-F**, n= 6) independent experiments. Statistical analysis: paired t-test (**A-C**), Mann–Whitney (**D-F, H-I**), Chi^2^ (**K**). ** p < 0.01; *** p < 0.001 ; **** p < 0.0001

Turnover of damaged mitochondria is part of the mitochondrial quality control process and it has recently been shown that enhanced mitophagy can slow-down the cell cycle progression (21). To measure mitochondria turnover in IL-7-treated WT and AMPK-KO Th cell cultures, we pulsed-chased mitochondria with the Mito-Green Tracker probe and followed the Mito-Green loss over time. We chose to perform these experiments on non-dividing IL-7-treated naive Th cells to avoid interferences between mitochondria turnover (on a per cell basis) and mitochondria dilution over cell divisions. As a control, cells were re-stained with the mitochondrial probe at each timing to check for total mitochondrial content. As shown in Figure 6A, B, Mito-Green staining gradually decreased over time and AMPK-KO Th cells exhibited increased mitochondrial turnover at each time point.

**Figure 6:**
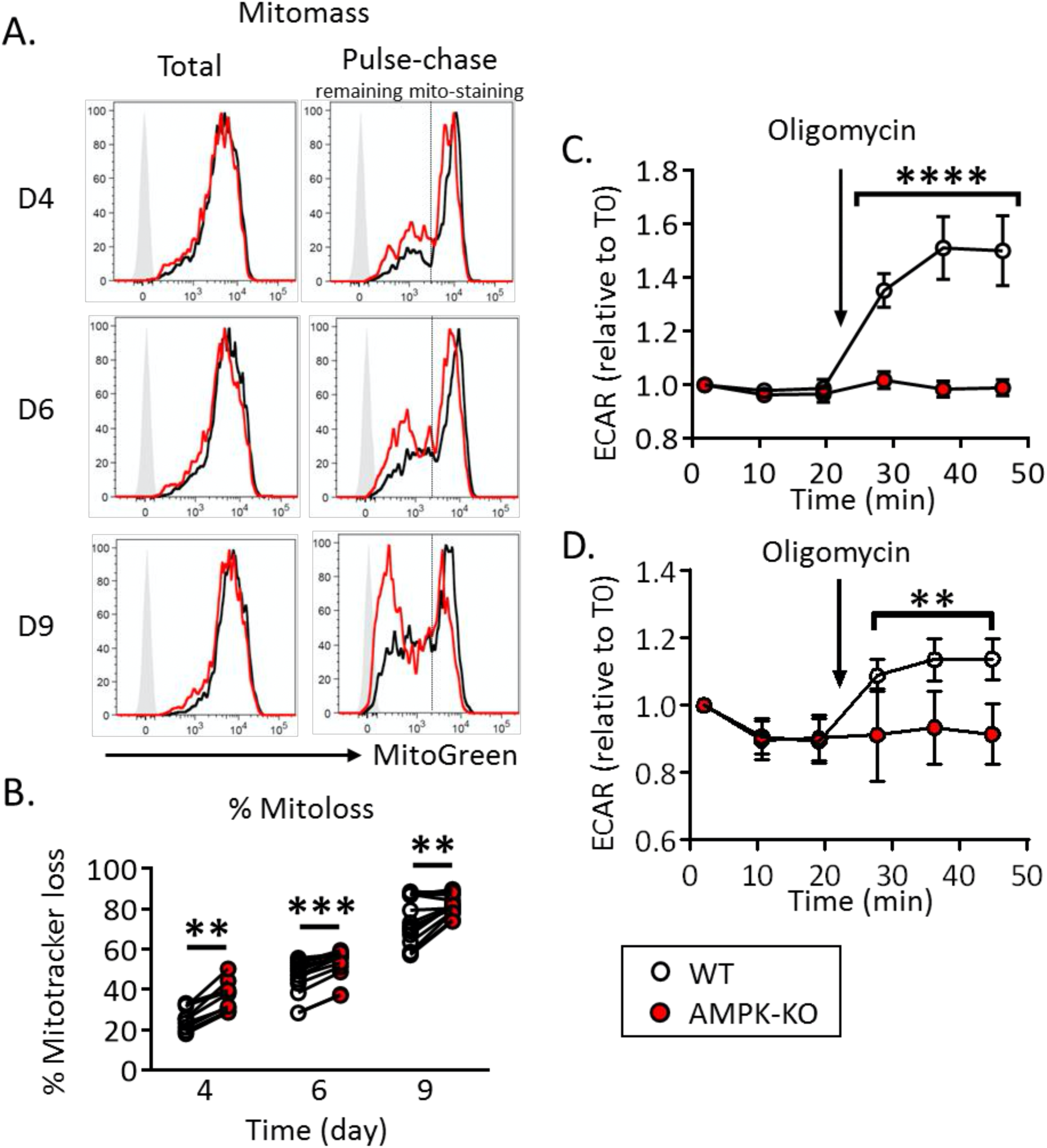
Accelerated mitochondrial clearance and defective mitochondrial stress-induced glycolytic switch in AMPK-KO T cells. (**A**)Representative FACS profiles of total mitochondrial mass (left panel) and pulse-chase remaining mitochondrial staining (right panels) in WT and AMPK-KO Th cells after 4, 6 and 9 day in IL-7-supplemented media. (**B**) Kinetic of mitochondrial mass loss, calculated from gate indicated in (**A**, right panels). (**C, D**) Kinetic of ECAR upon oligomycin treatment of WT and AMPK-KO Th1 memory-like cells (**C**) or IL-7+DC-expanded T cells (**D**). Data are pooled from 3 independent experiments (n = 9-16, **B**) or 2 independent experiments (n = 6, **D**), or are representative of 7 individual experiments (n = 16, **C**). Statistical analysis: Wilcoxon (**B**), Mann-Withney (**C, D**). ** p < 0.01 ; **** p < 0.0001

Mitochondrial fitness is essential for the rapid induction of glycolysis during re-activation of memory CD8^+^ T cells (22). Interestingly, WT but not AMPK-KO Th cells recovered from long term *in vitro* cultures showed increased ECAR upon both mitochondria stress and anti-CD3/CD28 re-stimulation (Figure 6C, D and supplemental Figure S10C).

### Dampening ROS production partially restores the competitive proliferative capacity of AMPK-KO T cells

Mitochondria are major cellular sources of reactive oxygen species (ROS) (23, 24). In agreement with a role for AMPK in maintaining redox balance in cancer cells (25, 26), AMPK-KO homeostatic proliferating T cells and memory-like Th1 cells generated increased amounts of ROS (Figure 7B, C). Although freshly purified or IL-7 cultured AMPK-KO naive Th cells expressed similar levels of intra-cellular ROS at steady state (Figure 7A and data not shown), they were highly sensitive to ROS-mediated cell death (Figure 7D), further indicating that AMPK regulates ROS balance and protects Th cells from ROS-mediated toxicity. WT and AMPK-KO Th cells exhibited similar sensitivity to a pro-apoptotic drug staurosporine (Figure 7 E), confirming the selectivity of AMPK for ROS-driven cell death. Similar results were obtained with memory Th1-like cells (Supplemental Figure S11).

**Figure 7:**
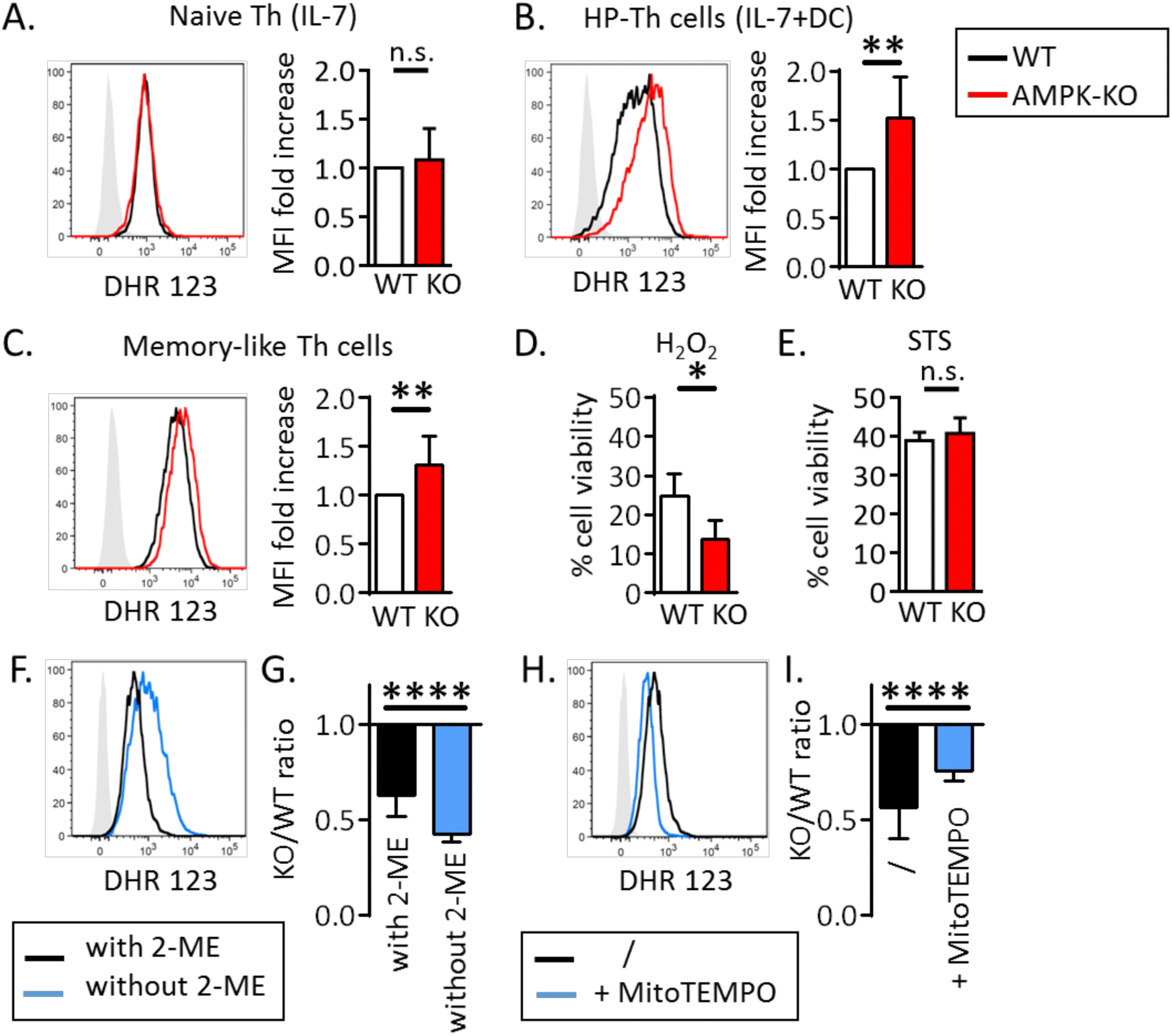
AMPK dampens ROS production and toxicity in T cells. (**A-C**) Cellular ROS staining with DHR123 probe in naive T cells (**A**), in in IL-7+DC-expanded T cells (**B**), and in memory-like Th1 cells (**C**). FACS staining of a representative experiments (left panels) and histogram showing MFI fold increase in AMPK-KO cells in comparison to WT cells set to 1 (right panels). (**D, E**) Percentages of cell viability after treatement of WT and AMPK-KO naive Th cells with H_2_O_2_ (0.5 mM for 2h, **D**) or with staurosporine (STS, 5 µM for 4h, **E**). (**F, H**) Cellular ROS staining of naive Th cells after 4 days in IL-7-supplemented culture medium with or without 2-ME (**F**) and supplemented with MitoTEMPO (**H**). (**G, I**) Relative recovery of AMPK-KO T cells after 14 days of homeostatic proliferation *in vitro* in a culture medium with or without 2-ME (**G**) and with or without MitoTEMPO (**I**). Data are pooled from 7 (n = 7, **A**), 9 (n = 11, **B**), (n = 8, **C**), 3 (n = 5, **D**), 4 (n = 4, **E**), 4 (n = 20, **G**) or 6 (n = 13, **I**) independent experiments or are representative of 3 individal experiments (**F, H**). Statistical analysis: Wilcoxon (**A-C**), Mann–Whitney (**D, E, G, I**). * p < 0.05; ** p < 0,01 ; **** p < 0,0001

Whereas low concentrations of ROS are essential for T cell activation and proliferation, high concentration of ROS affect cell survival, thereby highlighting a major role for anti-oxidative processes in the tight control of intra-cellular ROS equilibrium and cell proliferation (27). To test whether increased ROS levels expressed by AMPK-KO Th cells would impede their competitive proliferation, we set up homeostatic proliferation cultures in the absence or presence of antioxidants. In agreement with our working hypothesis, lack of 2-mercaptoethanol (2-ME), a potent reducing agent that prevents toxic levels of oxygen radicals through stabilization of the intracellular cysteine and GSH levels (28, 29), led to a reduced proliferative capacity of AMPK-KO when compared to WT cells (Figure 7F, G). Accordingly, supplementation of the culture media with the ROS scavenger MitoTEMPO reduced intra-cellular ROS contents and partially restored the proliferative competence of AMPK-KO Th cells (Figure 7H, I).

## Discussion

AMPK plays important roles in cellular energy metabolism and orchestrates cellular antioxidant defenses, thereby acting as a safeguard against prolonged and potently detrimental oxidative damages (25). We show herein that AMPK dampens intra-cellular ROS levels in homeostatic proliferating T cells and protects Th cells from ROS-mediated cell mortality, thereby sustaining prolonged proliferation both *in vitro* and *in vivo*. Moreover, during the course of GVHD, AMPK expression in alloreactive T cells drives T cell expansion and accelerates the onset of the disease. Because hematopoietic stem cell transplantation is likely associated with both beneficial (Graft versus Tumor) and pathological (Graft versus Host Disease) side effects (19, 30), understanding the mechanisms regulating the chronic T cell proliferation in response to continuously present alloantigens is of great importance.

Reactive oxygen species are generated as by-products of mitochondrial metabolism and eliminated via antioxidant systems. It is well established that, upon antigenic stimulation, T cells produce mitochondrial ROS that regulate secondary signaling molecules to bring about optimal TCR activation (31, 32). However, excessive mitochondrial ROS production or defective ROS neutralization by antioxidant cellular systems can also lead to T cell death, and antioxidant reagents can improve the survival of activated T cells (33–36). Cellular ROS levels increase with enhanced mitochondrial metabolic activity but are also produced under conditions of metabolic stress, such as hypoxia (37, 38).

In agreement with the well-known role of AMPK in regulating mitochondrial fitness (39), we observed that AMPK-deficient T cells displayed mitochondrial depolarization and accumulated mitochondrial membrane damages. Although AMPK has been reported to promote mitochondrial biogenesis through activation of the transcription factor PGC1 in distinct cell types (39), AMPK-KO T cells displayed similar mitochondrial mass and *pgc1* expression, indicating that the mitochondrial defects observed in this experimental model occurred independently of total mitochondrial cell content. *In vitro*-expanded AMPK-KO T cells exhibit a similar mitochondrial respiration rate, compared to their wild type counterparts. This contrasts with data showing that cytotoxic T effector cells isolated from AMPK-deficient mice infected with *Listeria monocytogenes* displayed reduced mitochondrial respiratory rates relative to control cells (17). Intriguingly, maximal OCR only slightly increased above basal OCR (low spare respiratory capacity, SRC), indicating that Th cell operate at their maximal oxidative rate in our proliferation settings. Use of SRC by naive T cells to energetically support *in vivo* spontaneous homeostatic proliferation has been recently reported (40). Utilization of SRC might thus increase oxygen consumption to maximal levels independently of mitochondrial potential. A sustained maximal respiration rate combined with defects in mitochondria membranes could therefore lead to increased ROS production in AMPK-KO proliferating T cells.

Freshly purified AMPK-KO naive T cells exhibited reduced mitochondrial potential without sign of excessive cellular ROS concentration (data not shown), indicating that a mitochondrial defect preceded ROS production during proliferation of AMPK-deficient T lymphocytes.

Alternatively, a defect in ROS scavenging mechanisms could also contribute to the gradual accumulation of ROS in proliferating AMPK-KO T cells overtime. One of the most striking observations of this study is the inability of homeostatic-expanded AMPK-deficient Th cells to shift towards a glycolytic metabolism in response to a mitochondrial poison or TCR signaling. To date, the role of AMPK in the control of the Warburg effect has been controversial. Inactivation of AMPKα in both transformed and non-transformed cells promoted a HIF-1α-dependent reprogramming of glucose metabolism characteristic of the Warburg effect (41). In contrast, loss of the tumor suppressor folliculin (FLCN) gene, a negative regulator of AMPK, promoted constitutive AMPK activation in cancer cells, leading to increased mitochondrial biogenesis and a ROS-induced HIF transcriptional activity, thereby coupling AMPK-dependent mitochondrial biogenesis to HIF-dependent Warburg metabolic reprogramming (42). Moreover, other reports identified AMPK as a major driver of glycolysis either through direct phosphorylation of PFKFB3 (43, 44) or indirectly, through mitochondrial ATP-driven hexokinase activation (22).

Our study thus posits AMPK as a driver of the Warburg effect in T lymphocytes. Whether the inability of AMPK-deficient T cells to switch back to glycolysis account for their defective proliferation awaits further experimentations.

In the absence of AMPK, homeostatic proliferating Th cells secreted reduced amount of IFNγ, compared to their wild type counterparts. This contrast with data showing that AMPK dampened IFNγ production in CD8^+^ T cells activated *in vitro*, in particular when cells are exposed to low glucose concentration (17). This discrepancy could be easily explained by the differences in the experimental models between the studies. Reduced IFNγ production did not reflect a general defect in cytokine secretion as Th cells of both genotypes produced similar amounts of IL-2. During homeostatic proliferation, IFNγ was secreted by fast dividing cells. It is therefore possible that higher IFNγ secretion in the wild type cell compartment simply reflected the presence of cells that underwent more division in this group. Although we excluded slow-divided cells (0-3 cell divisions, based on CFSE contents) from the analysis, we could not test this hypothesis further as CFSE dilutions did not allow us to discriminate cells undergoing more divisions. Decreased IFNγ secretion by AMPK-KO T cells could also result from their defective glycolytic switch as IFNγ production was intimately connected with a glycolytic metabolism (22, 45, 46).

The observation that AMPK deficiency did not preclude Th cell proliferation in response to a strong mitogenic anti-CD3/CD28 signaling confirms and extends previous data showing that AMPK is dispensable for a short term effector cell differentiation (15, 16). A strong Akt / mTOR-driven glycolytic program might thus compensate for AMPK deficiency and sustain a rapid proliferative response (47). However, anti-CD3/CD28 activated AMPK-KO T cells exhibited defective proliferation when expanded with IL-2 for longer periods. Taken together, our data thus indicate that the homeostatic proliferative defect of AMPK-KO T cells does not reflect a specific defective response to homeostatic signals but rather results from the progressive accumulation of uncontrolled ROS production, leading to cell damages and premature senescence. In support of this assumption, we observed that antioxidant treatments partially alleviated the competitive proliferative defect of AMPK-KO T cells. In this context, an accelerated mitochondrial component turnover could reflect an increased recycling of damaged mitochondria in AMPK-KO T cells, although this hypothesis still warrants further investigations.

AMPK was recently shown to down-regulate PD-1 expression, thereby promoting memory-like activities of CAR-T cells *in vivo* (48). Of interest, PD-1 mediated regulation of T cell function through repressed mitochondrial metabolism and production of ROS, potentially leading to cell death of activated T cells (49, 50). A fine tuning of the AMPK mediated control over cell metabolism and ROS equilibrium would thus open new avenues for medical interventions in order to prevent GVHD or to sustain CAR-T cell homeostatic expansion *in vivo*.

## Materials and methods

### Mouse model

AMPKα1^fl/fl^ mice were kindly provided by Dr. B. Viollet (Institut Cochin, Paris - France) and bred to CD4-Cre transgenic mice, obtained from Dr. G. Van Loo (University of Gand - Belgium). C57BL/6 and Balb/c mice were purchased from Envigo (Horst - Nederland). CD45.1 were purchased from Jackson Laboratory (Bar Harbor - USA) and CD3ε^-/-^ were obtained from EMMA (Orléans – France). Control mice were littermate AMPKα1^fl/fl^ Cre^-^ or AMPKα1^+/fl^ Cre^-^ or sex- and age-matched C57BL/6 or CD45.1 mice. Mice were housed under specific pathogen-free (SPF) conditions. The experiments were performed in compliance with the relevant laws and institutional guidelines and were approved by the Local Ethic Committee.

### Cell purifications

Bone marrow cells from tibia and femur were purified from naive mice by flushing. When required (GVHD), T cells were magnetically depleted. Briefly, bone marrow cells were incubated 20 min at 4°C with anti-CD4 (L3T4)-, anti-CD8α (Ly-2)- and anti-CD90.2-MicroBeads (Miltenyi Biotec, Bergisch-Gladbach, Germany) and then separated over an auto-MACS column using the DepleteS program.

Total T cells (CD4^+^ and CD8^+^) were purified from the spleen of naive mice by Stemcell negative selection, according to manufacturer’s recommendations. Briefly, spleen cells were incubated 30 min at 4°C with 0,5 µg/ml of anti-Ter119-biot (Ly-76), anti-CD19-biot (MB19-1), anti-Gr1-biot (RB6-8C5), anti-CD49b-biot (DX5), anti-CD11c-biot (N418) and anti-MHCII (I-A/I-E)-biot (M5/114) antibodies. Antibodies are from BD Biosciences or BioXCell. Cells were then incubated with Biotin selection cocktail 2 (Stemcell) for 17 min at 4°C. D2 Magnetic Particles (Stemcell) were added and cells were incubated for an additional 7 min at 4°C before placing them in the Stemcell magnet.

Naive T cells (CD4^+^ CD44^-^) were purified from the spleen of naive mice by MACS column enrichment followed by cells sorting. Spleen cells were incubated at 4°C with anti-CD4 (L3T4)-MicroBeads (Miltenyi Biotec) after 10 min CD4 (RM4-5)-PB, CD44 (IM7)-PE-Cy7, CD62L (MEL-14)-A700 and CD25 (PC61)-BB515 were added for an additional 15 min. These antibodies are all from BD. The cells were first separated over an LS-column (Miltenyi Biotec) and the CD4^+^ fraction was then sorted as CD4^+^ CD25^-^ CD62L^+^ CD44^-^ by FACS ARIA III (BD Biosciences).

For dendritic cell purification, spleens were treated with collagenase (Gestimed). Recovered splenocytes were resuspended in Nicodenz (DO_360_=0.666, Lucron Bioproducts). RPMI-medium with 3% FCS was added on the top of Nicodenz. After 15 min centrifugation at 4°C and 1700G without brake, DC fraction were recovered from the gradient interface and incubated with CD11c-MicroBeads (Miltenyi Biotec) for 15 min at 4°C. The excess of beads was washed out and cells were then separated over an LS-column (Miltenyi Biotec).

### Bone marrow transfer

Bone marrow cells purified from WT (CD45.1) and AMPK^KO-T^ (CD45.2) mice were mixed at ratio 1:1 and i.v. injected (4 to 5.10^6^ cells in total) into 1200 cGy (split dose) irradiated CD3ε^-/-^ mice. 8 to 12 weeks later, WT and AMPK-KO T cells from different organs were analyzed by flow cytometry.

### *In vivo* homeostatic proliferation

Naive (CD4^+^ CD44^-^) cells obtained from WT (CD45.1) and WT or AMPK^KO-T^ (CD45.2) mice were mixed at ratio 1:1 and i.v. injected (1-2.10^6^ cells in total) into CD3ε^-/-^ mice. 10 to 12 days later, WT and AMPK-KO T cells from different organs are analyzed by flow cytometry. In some experiments cells were CFSE-labelled (Invitrogen) before i.v. injection.

### GVHD

Irradiated Balb/c mice (750 cGy, day -1) were i.v. injected with 5.10^6^ C57BL/6 T cell depleted bone marrow with or without 10^6^ total T cells from either WT or AMPK^KO-T^ mice (day 0). Clinical score was followed three times a week and calculated as previously described by Cooke et al, 1996 (51). 7 to 10 days later, WT and AMPK-KO T cells from different organs are analyzed by flow cytometry.

### Isolation of intestinal lymphocytes

Intestines were collected and cleared from residual connective tissue, Peyer’s patches and feces. Intestines were then cut into small pieces and incubated for 30 min at 37°C with agitation in a medium containing EDTA (2.5 mM, VWR) and DTT (72.5 µg/ml, Sigma-Aldrich). IEL were recovered by washing and shaking the intestines 3 times in a medium with EDTA and one time without EDTA. To obtain LP lymphocytes, intestines were further chopped and incubated for 30 min at 37°C in medium containing liberase (20 µg/ml, Roche) and DNAse (400 µg/ml, Roche). Enzymes were neutralized with EDTA (2.5 mM) and the intestines were crushed and filtered. IEL and LP fractions were finally washed and submitted to 15 min centrifugation at 500G, 20°C without break in a 30% Percoll (GE Healthcare) solution. Lymphocytes are in the pellet.

### *In vitro* differentiation of memory-like Th cells

Naive T cells (CD4^+^ CD44^-^) were polarized into Th1 cells for 3 days using plate-bound anti-CD3 (5 µg/ml, clone 145-2C11, BioXCell), anti-CD28 (1 µg/ml, clone 37.51, BioXCell), recombinant IL-12 (10 ng/ml, Peprotech) and anti-IL-4 (10 µg/ml, BioXCell). Th1 cells were then cultured, for 4 to 7 additional days, into IL-7-supplemented (20 ng/ml, Peprotech) media to obtain memory-like cells.

### *In vitro* homeostatic proliferation

Naive T cells (CD4^+^ CD44^-^) purified from WT (CD45.1) and AMPK^KO-T^ (CD45.2) mice were mixed at a 1:1 ratio, CFSE-labelled and seeded in a 96-well plate (10^5^ cell/well). T cells were cultured at 37°C in RPMI supplemented with 5% heat-inactivated FBS (Sigma-Aldrich), 1% non-essential amino acids (Invitrogen), 1 mM sodium pyruvate (Invitrogen), 2 mM L-glutamin (Invitrogen), 500 U/mL penicillin/500 μg/ml streptomycin (Invitrogen), and 50 μM β-mercaptoethanol (Sigma-Aldrich). The homeostatic stimuli were provided by 2.10^4^ DC purified from autologous C57BL/6 (H-2^b^) mice and IL-7 (20 ng/ml, Peprotech). Each 4-5 days of culture, half of the medium volume was refreshed with new IL-7-supplemented medium and freshly purified DC. Cells were analyzed on day 13-15. For antioxidant supplementation, MitoTEMPO (20 µM, Sigma) was added to culture medium and refreshed at the same time than IL-7.

Memory-like Th1 from WT (CD45.1) and AMPK^KO-T^ (CD45.2) mice were mixed at ratio 1.1 and were cultured with 2.10^4^ syngeneic DC and IL-7 (20 ng/ml, Peprotech) for 9 additional days. Half of the medium volume was refreshed with new IL-7-supplemented medium and freshly purified DC at day 5.

### Flow cytometry

Specific cell-surface staining was performed using standard procedure with the following monoclonal antibodies conjugated to fluorochrome or biotin: CD4 (RM4-5), CD44 (IM7), CD45.1 (A20) from BD Bioscience, and CD8α (53-6.7), CD183 (CXCR3-173), CD62L (MEL-

14), TCRβ (H57-597) from eBioscience. Strepatividin was purchased from BD Bioscience. Dead cells were excluded by LIVE/DEAD (Invitrogen) staining and non-specific, Fc-mediated interactions were blocked by anti-Mouse CD16/CD32 (2.4G2 BioXCell). For intracellular staining, cells were incubated with a fixing solution (Fixation/Permeabilization, eBioscience) and stained with directly conjugated monoclonal antibodies anti-IFNγ (XMG1.2), IL-2 (JES6-5H4), Ki67 (B56) from BD-Bioscience, and FoxP3 (FJK-16s) from eBioscience. For cytokine production assessment, cells were pre-incubated for 4 h with PMA 50 ng/ml (Sigma), ionomycin 1 µg/ml (Sigma) and brefeldin A (eBiosciences).

For mitochondrial parameters analysis, cells were incubated 25 min at 37°C with the Mito-Green dye (100 nM, Invitrogen), TMRE (5 nM, Invitrogen) or JC-1 probe (2 µM, Invitrogen) in complete medium.

For the mitochondria renewal assay, naive T cells were stained with Mito-Green dye (2 µM, Invitrogen) for 20 min at 37°C (10^6^ cells per 100 µL). Then the loss of this staining was recorded by flow cytometry at day 4, 6 and 9. At each time point, fresh staining of Mito-Green dye was performed to measure total mitochondria content on unstained cells (100 nM). For ROS detection, cells were incubated 30 min at 37°C with Dihydrorhodamine (DHR) 123 (2 µM, Sigma) in complete medium. Cells were analyzed by FACS Canto II or ARIA III (BD Bioscience) and data were analyzed with the FlowJo Software (Tree Star).

### Seahorse

OCR and ECAR were determined using a Seahorse XFp or XF96 Extracellular Flux Analyser following user guide. Plate were pre-coated with poly-D-lysine (200 µg/ml, Millipore) for cell adhesion. 1 mM pyruvate, 2 mM de glutamine and 10 mM glucose were added to XF Base Medium and pH was adjusted to 7.4. Drugs were added to reach a final concentration of: oligomycin 1 µM, FCCP 0.5 µM, rotenone/antimycin A 0.5µM and 2-DG 50mM. For CD3/CD28 beads (Dynabeads Mouse T-Activator CD3/CD28, LifeTechnologies) injection, beads:cells ratio was 4:1. Seahorse protocol was adjusted with the following settings: mix 2min, wait 2min and measure 4-5min. At the end of the Seahorse analysis, cells were lysed and Bradford (Biorad) protein dosage was performed.

### ROS / staurosporine-sensitivity

2-5.10^5^ cells were incubated for 2 h with different concentrations of H_2_O_2_ (Merck) or for 2, 4 or 6 h with staurosporine (5 µg/ml, Sigma) at 37°C. The percentage of survival was measured after Propidium iodide staining (Sigma).

### Transmission Electron Microscopy (TEM)

A total of 27 (WT) and 29 (AMPK-KO) T cells were analyzed by transmission electron microscopy. Naive T cells were maintained 4 days in IL-7-supplemented medium and then fixed overnight in 2.5% glutaraldehyde (Electron Microscopy Sciences, UK) at 4°C and post-fixed with 1% osmium tetroxide (Electron Microscopy Sciences, UK) and 1,5 % ferrocyanide (Electron Microscopy Sciences, UK) in 0,15 M cacodylate buffer. The cells were further stained with 1% uranyl acetate (Electron Microscopy Sciences, UK), serially dehydrated and embedded in epoxy resin (Agar 100 resin; Agar Scientific, UK). Ultrathin 70-nm sections were produced with a Leica EM UC6 ultramicrotome and mounted on copper–Formvar–carbon grids (Electron Microscopy Sciences). Observations were made on a Tecnai 10 electron microscope (FEI, Eindhoven, Netherlands), and images were captured with a Veleta camera and processed with SIS iTEM sofware (Olympus, Germany).

## Statistical analysis

Statistical analysis were performed using Prism6 (GraphPad Software, San Diego, CA). Datasets were first tested for Gaussian distribution with the D’Agostino & Pearson omnibus normality test. Statistical analysis performed are detailed for each graph in the figure legend. n.s.: non significant; *p < 0,05; **p< 0,01; ***p< 0,001; ****p< 0,0001.

## Aknowledgments

We especially thank Muriel Moser for her great support all along this work, for stimulating discussions and for careful revision of the manuscript. We also thank Caroline Abdelaziz and Véronique Dissy for animal care and for technical support. This work was supported by the European Regional Development Fund (ERDF) and the Walloon Region (Wallonia-Biomed portfolio, 411132-957270), grant from the Fonds Jean Brachet and research credit from the National Fund for Scientific Research, FNRS, Belgium. The CMMI is supported by the European Regional Development Fund and the Walloon Region.

F.A. is a Research Associate at the FNRS. T.P. is recipient of a research fellowship from the FNRS/Télévie. A.L. is supported by a Belgian FRIA fellowship.

## Figure legends

**Figure S1:**
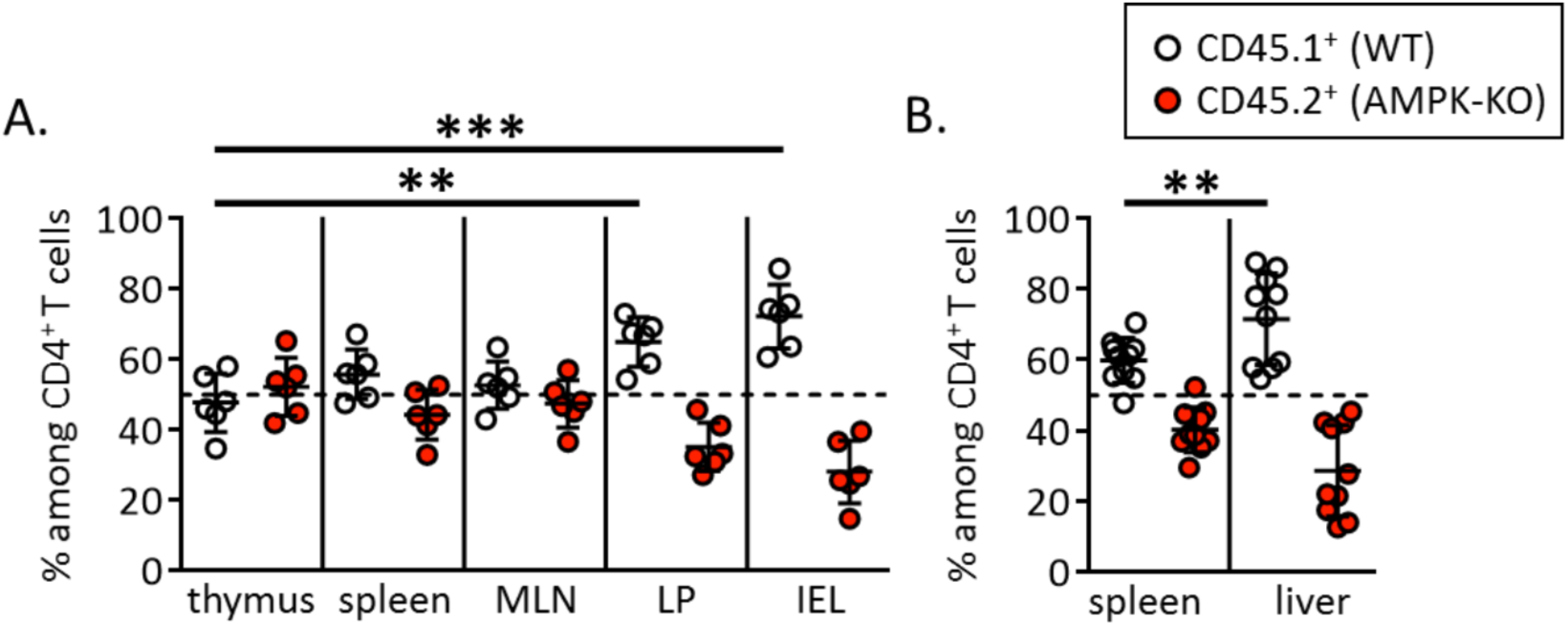
AMPK promotes peripheral T cells replenishment upon bone marrow transfer. **(A-B)** Relative contribution of AMPK-KO CD4^+^ to the repopulation of the thymus, spleen, M LN, LP and IEL **(A)** of CD3ε^-/-^ mice were reconstituted with a 1:1 mixture of bone marrow cells from WT (CD45.l) and AMPK^KO-T^ (CD45.2) mice as in Figure 1 (symbols represent individual mice; 1 experiment representative of 3 with n = 6) and **(B)** spleen and liver (pool of 2 independent experiments with n= 10). Statistical analysis: Friedman followed by Dunn multiple comparison **(A)**, Wilcoxon **(B).** ** p < 0.01; *** p < 0.001 MLN = Mesenteric lymph nodes, LP= Lamina propria, IEL = lntraepithelial lymphocytes

**Figure S2:**
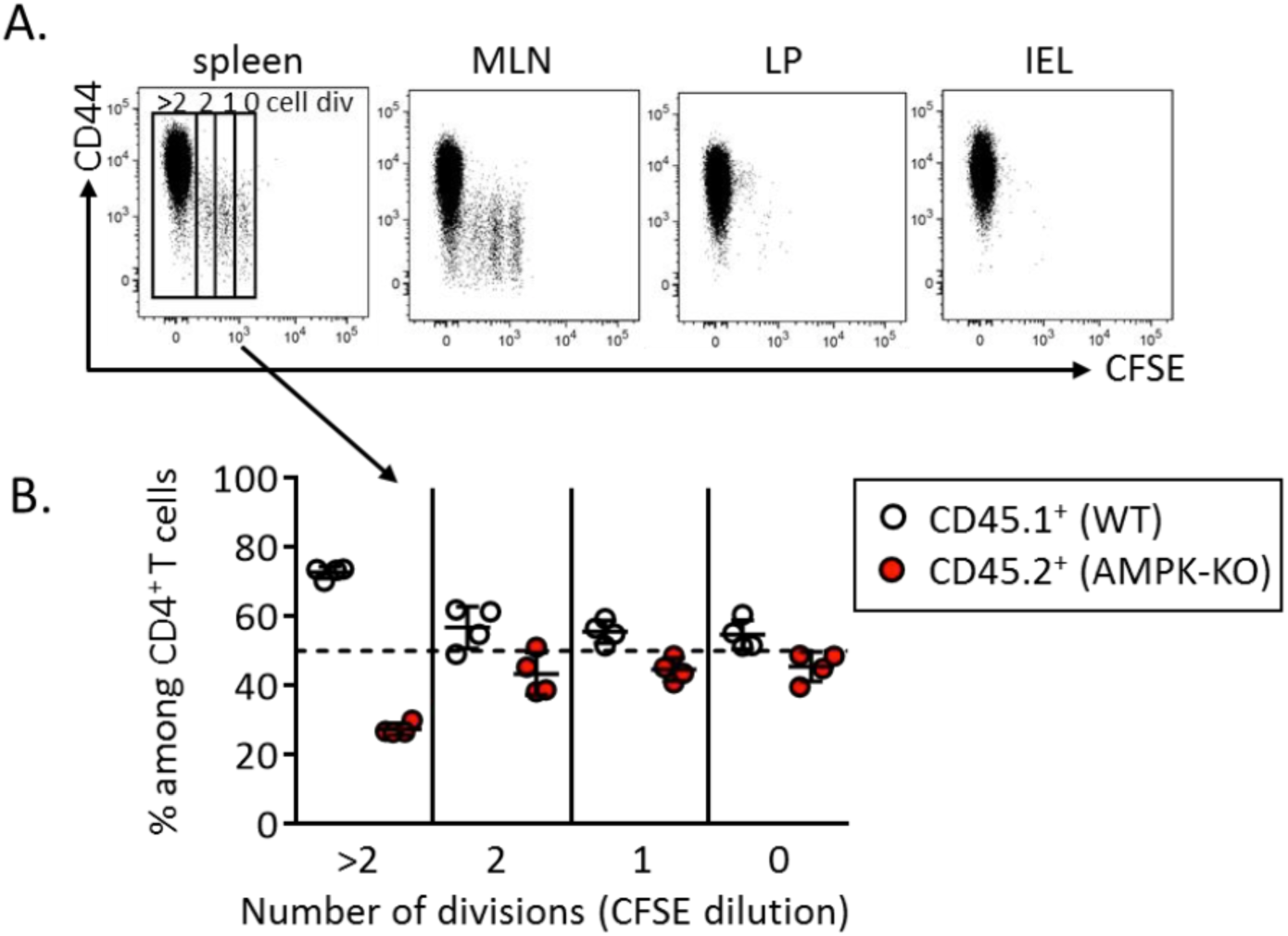
AMPK gradually promotes accumulation of divided cells during competitive homeostatic proliferation. CFSE-labeled WT (CD45.l) and AMPK ^KO-T^ (CD45.2) naive T cell (1:1 ratio) were i.v. injected into CD3ε^-/-^ mice. Recovered WT and AMPK-KO T cells were analyzed by flow cytometry 10 days later. **(A)** Expression of CD44 according to the number of cell divisions (CFSE dilution) in the spleen, MLN, LP and IEL of recipient mice. **(B)** Relative contribution of WT and AMPK-KO CD4^+^ T cells to the repopulation of the spleen according to the number of cell divisions (1 experiment representative of 3 with n= 4). MLN = Mesenteric lymph nodes, LP= Lamina propria, IEL = lntraepithelial lymphocytes

**Figure S3:**
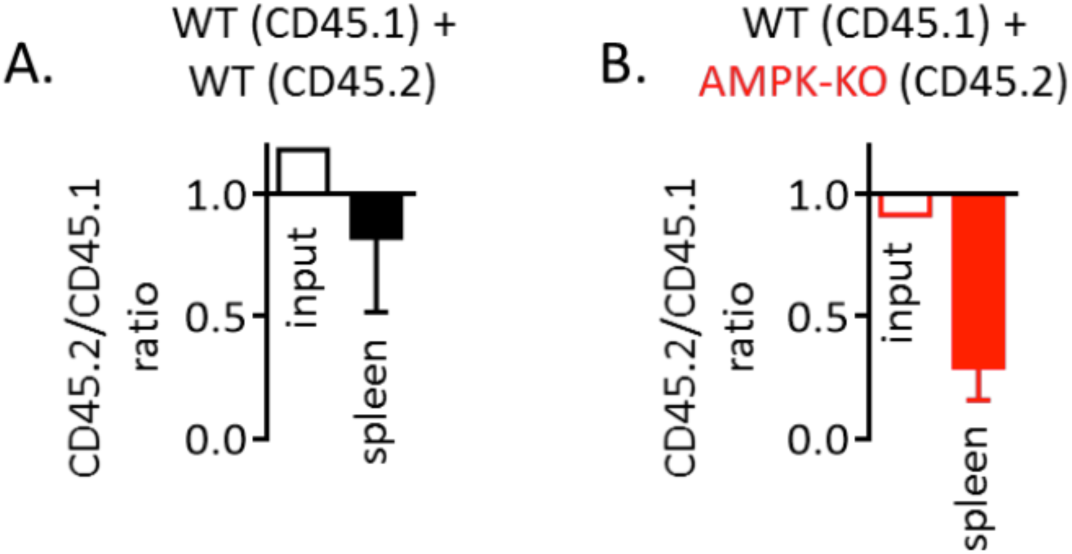
AMPK-KO T cell defect in homeostatic proliferation does not result from CD45 polymorphism. WT (CD45.1) naive T cells were mixed with mice WT **(A)** or AMPK-KO **(B)** counterparts (both from CD45.2 mice) and inoculated into CD3 ε^-/-^ mice. Results represent the relative contribution of CD45.2 cells in the inoculum (input) and in the spleen of recipient mice on day 10 after transfer (mean +/- SD of 3 recipient mice).

**Figure S4:**
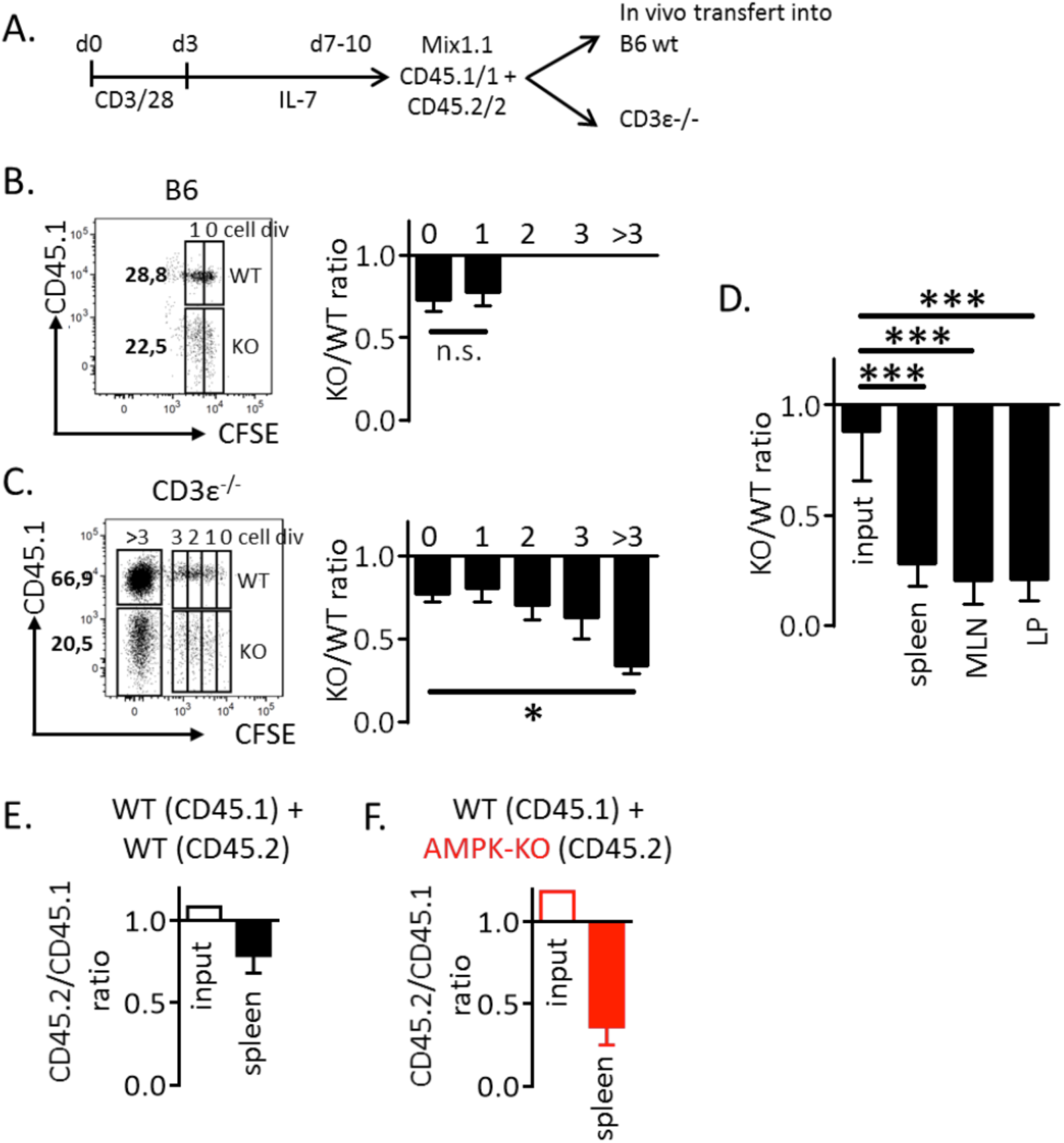
AMPK-KO memory-like Th cells exhibit impaired homeostatic proliferation. **(A-D)** IL-7-driven memory-like Th1 cells from WT (CD45.1) and AMPK ^KO-T^ (CD45.2) mice were mixed at 1:1 ratio and injected i.v. into C57BL/6 **(B)** or CD3 ε^-/-^ **(C-D)** mice. Recovered T cells were analyzed on day 10 by flow cytometry. **(A)** Experimental protocol. **(B, C)** Dot plots and histograms show the relative proportions of WT and AMPK-KO cells in the spleen of recipient mice (CD4^+^ gated), according to CFSE division status. Numbers upper each box indicate cell division number, numbers on the left side indicate the percentage of cells in the corresponding gate. **(D)** Relative contribution of AMPK-KO memory Th1 cells to the repopulation of the CD4^+^ T cell subset in the spleen, MLN and LP of CD3ε^-/-^ mice. Data are representative of 3 independent experiments with n = 5 **(B, C**) or pooled from 3 experimenst, with n = 13 **(D)**. **(E-F)** Memory-like Th1 cells from WT (CD45.1) mice were mixed with WT **(E)** or AMPK-KO **(F)** (both CD45.2) counterparts and inoculated into CD3ε^-/-^ mice. Results represent the relative contribution of CD45.2 cells in the inoculum (input) and in the spleen of recipient mice on day 10 after transfer (mean +/- SD of 3 recipient mice). Statistical analysis: Wilcoxon **(B, D)**, Friedman followed by Dunn multiple comparison **(C)**. * p < 0,05, *** p < 0,001 MLN = Mesenteric lymph nodes, LP= Lamina propria

**Figure S5:**
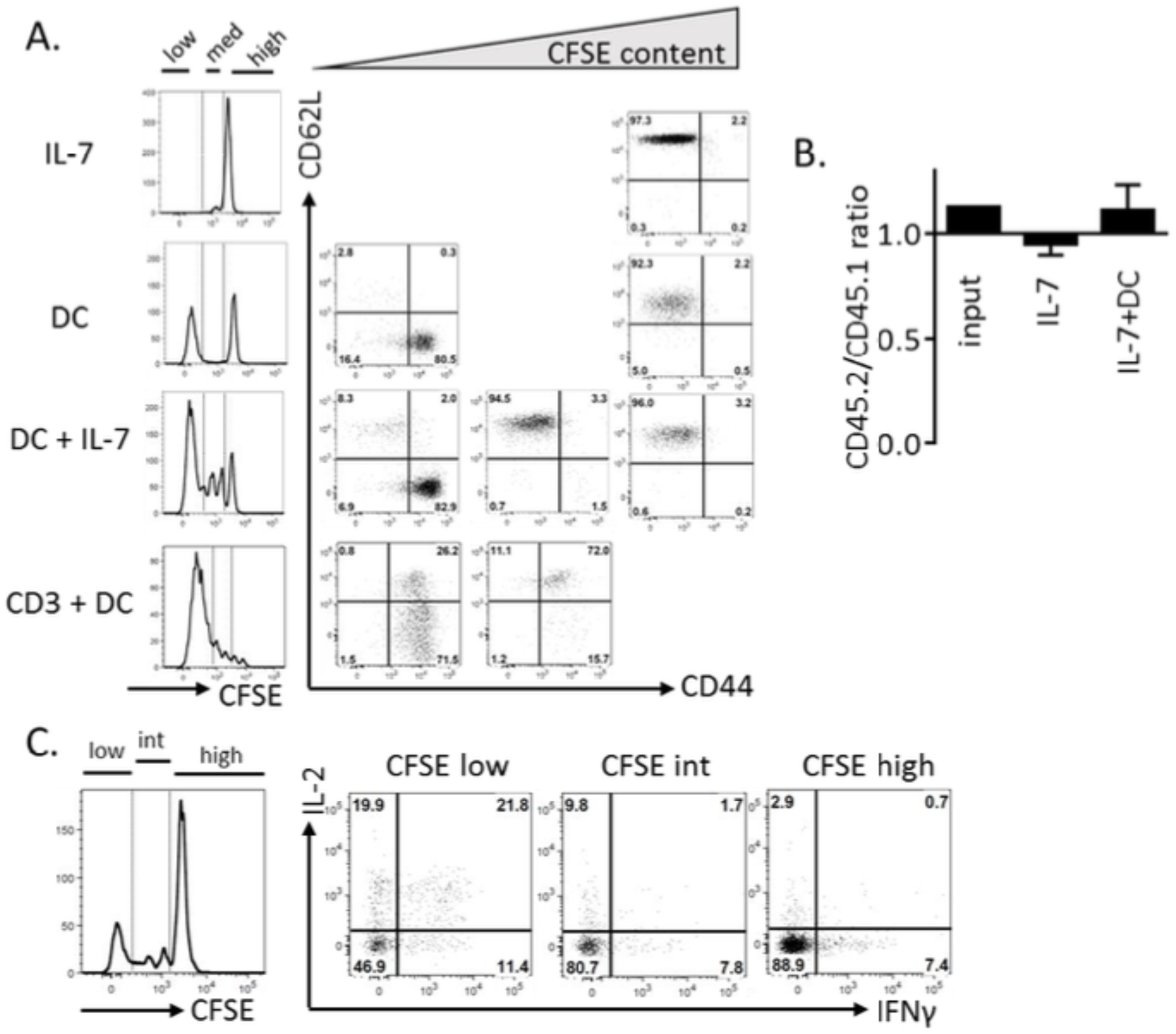
Model of IL-7 driven homeostatic proliferation *in vitro*. **(A)** Naive T cells from CS7BL/6 mice were labeled with CFSE and cultured in the presence of IL-7 and/ or syngeneic DCs, as indicated. Culture media were replaced with fresh DCs and IL-7 each 4-5 day and cells were analyzed on day 14 and 18. Additional control included cells activated in the presence of DCs + anti-CD3 mAbs for 5 days. CFSE dilution (left panels) and CD44 / CD62L expression (gate CD4^+^) as a function of CFSE content. **(B)** Naïve T cells from WT (CD45.1) mice were mixed with WT (CD45.2) counterparts and cultured in the presence of IL-7 (n=8) or IL-7+syngeneic DCs (n=12). Results represent the relative contribution of CD45.2 cells in the inoculum (input) and after 13 days of culture. **(C)** Naive T cells from CS7BL/6 mice were labeled with CFSE and cultured in the presence of IL-7 and syngeneic DCs. Percentage of CD4^+^ T cells producing IL-2 and IFNγ according to the number of cell divisions after 14 days of culture.

**Figure S6:**
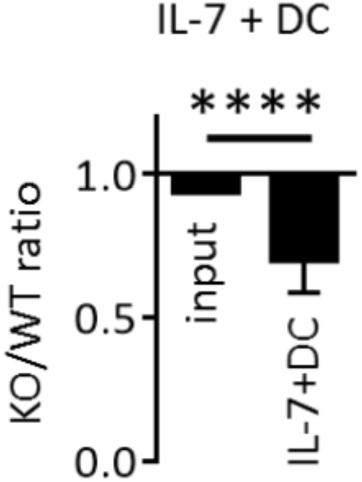
AMPK-KO memory-like Th cells exhibit impaired homeostatic proliferation *in vitro*. Relative recovery of AMPK-KO memory-like Th1 cells after a 9 day-homeostatic culture with IL-7 and syngeneic DCs. Statistical analysis: Wilcoxon;**** p < 0.0001

**Figure S7:**
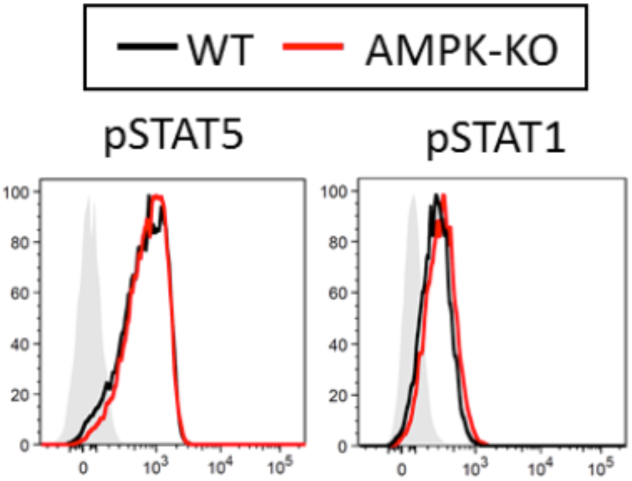
AMPK-KO Th cell express normal phospho-STAT5/STAT1 signaling in response to IL-7. Phosphorylation levels of STAT5 and STAT1 after 30 min of IL-7 treatment (data are representative of at least 3 experiments). Filled grey histograms represent unstained Th cells.

**Figure S8:**
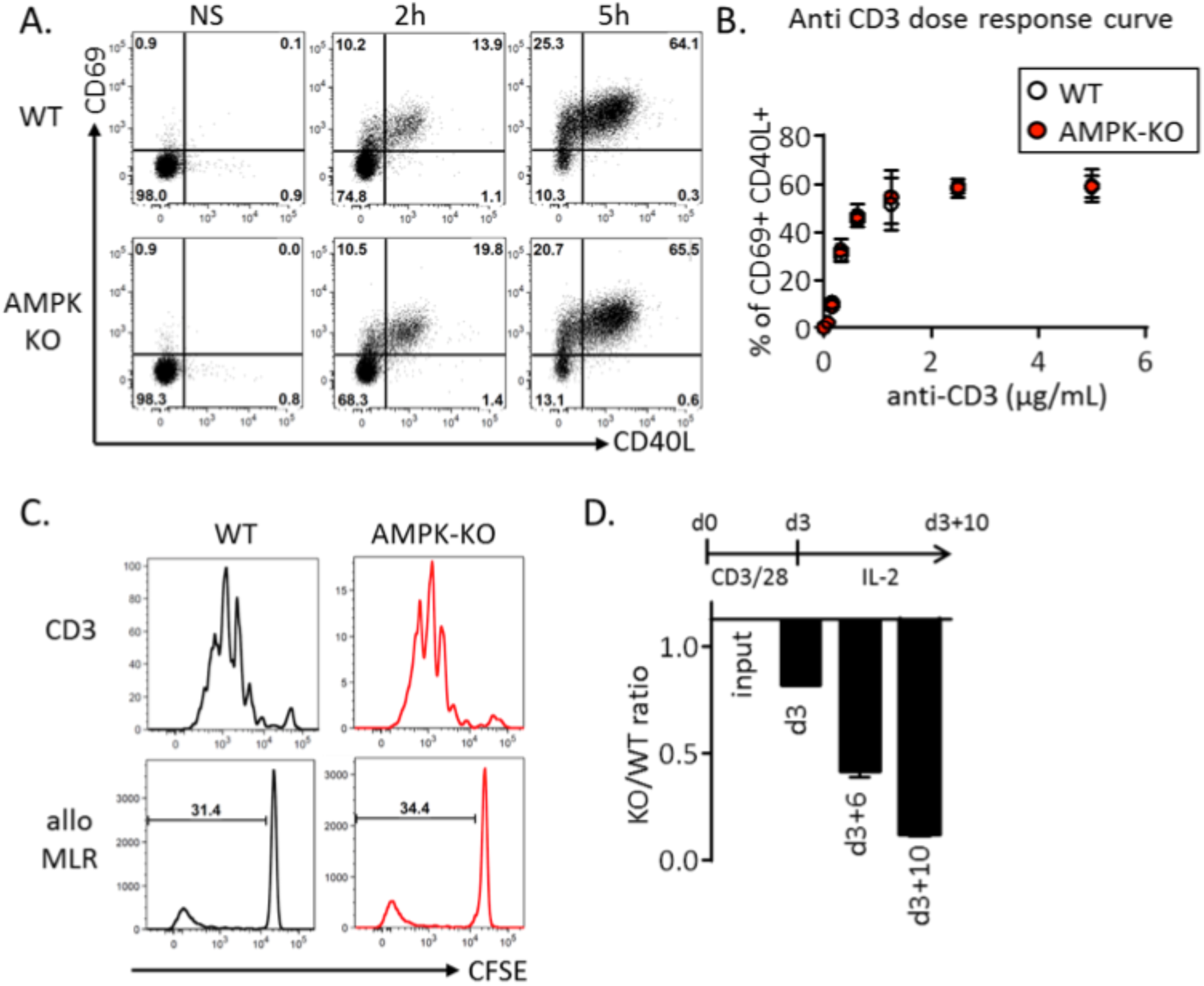
AMPK is not required for early naive T cell activation and proliferation but confers a competitive advantage during sustained T cell expansion. **(A)** Early acquisition of the activation markers CD69 (H1.2F3, eBioscience) and CD40L (MR1, eBioscience) by WT and AMPK-KO memory-like Th cells after 2 and 5h of anti-CD3/ CD28 stimulation (1 experiment representative of 3). **(B)** Dose response to anti-CD3 stimulation of WT and AMPK-KO memory-like Th cells after 2h (pool of 2 independent experiments, n=2). **(C)** CFSE dilution profiles of WT and AMPK-KO naive T cells stimulated with anti-CD3 / CD28 mAbs (upper panel) or Balb/c-derived allogeneic DCs (lower panel) during 5 days (1 experiment representative of 3). **(D)** Naive T cells were polarized into Th1 cells for 3 days with plate-bound anti-CD3, anti-CD28, IL-12 and anti -IL-4. Th1 cells were further diluted and cultured for an additional 10 days in IL-2-supplemented (20 ng/ml, Peprotech) media to obtain effector-like cells. Data express the relative recovery of AMPK-KO T cells after anti-CD3/ CD28 stimulation (day 3) and after 6 and10 day of IL-2-supplemented culture (day 3+6 and 3+10, respectively). 1 experiment representative of 2, with n = 2 (day 3) or n = 5 (day 3+6 and day 3+10). NS = unstimulated

**Figure S9:**
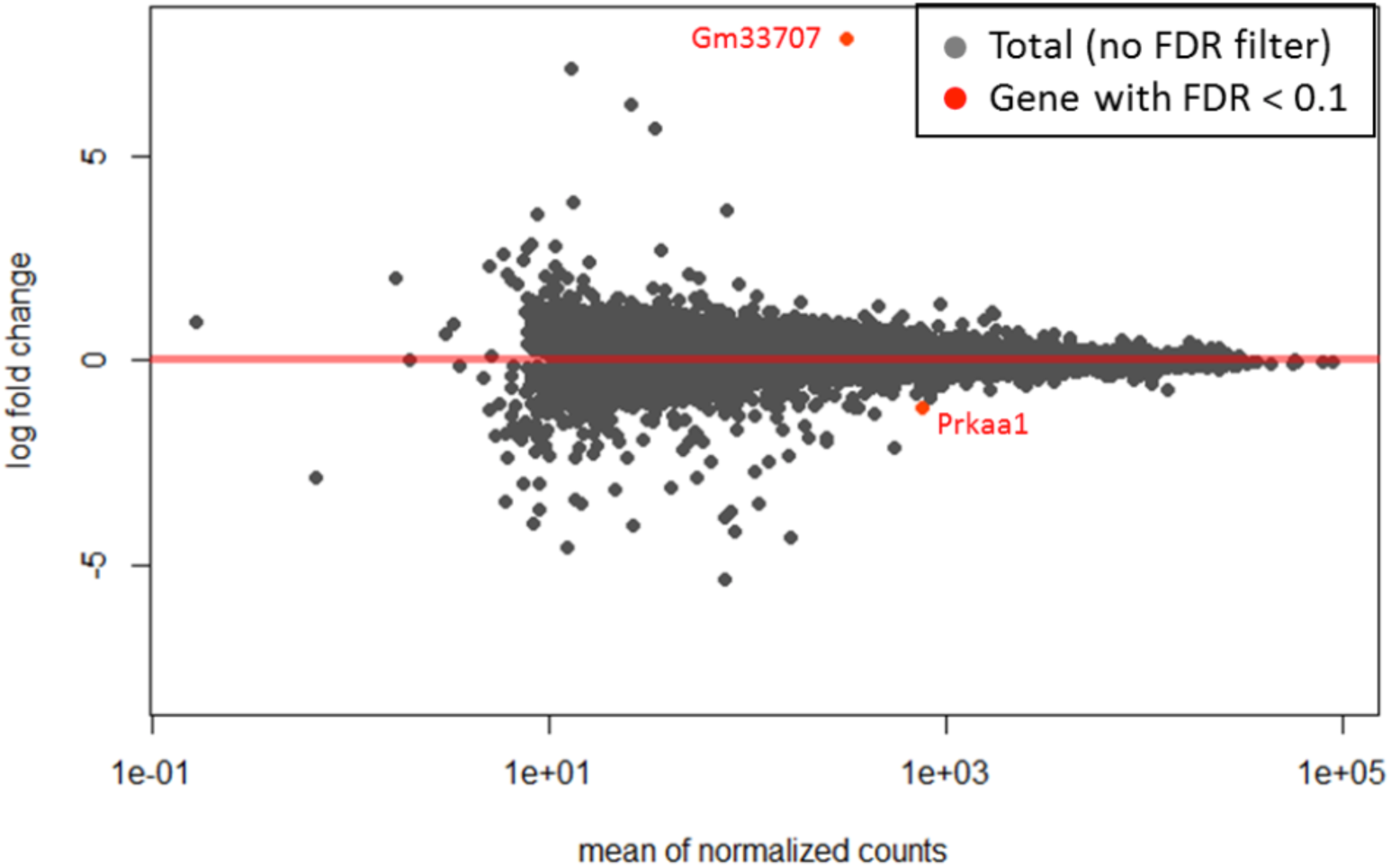
Homeostatic proliferating WT and AMPK-KO T helper cells express a similar gene pattern. WT and AMPK-KO naïve T cells were recovered after of 14 days of HP with IL-7 and syngeneic DCs and subjected to RNAseq analysis (1 experiment, n = 3 per group). **(A)** MA of the differentially expressed genes (DEGs) with low expression genes (less than 20 read in WT) excluded. Y-axis: logFC (fold change); X-axis: average expression for each gene; the color of the data points denotes the status of DEGs (red points: FDR <0.1; grey points: with FDR >0.1). CPM, counts per million; FDR, false discovery rate. Only two transcripts: Gm33707 and Prkaa1 show distinct expression between WT and AMPK-KO (FDR< 0.1).

**Figure S10:**
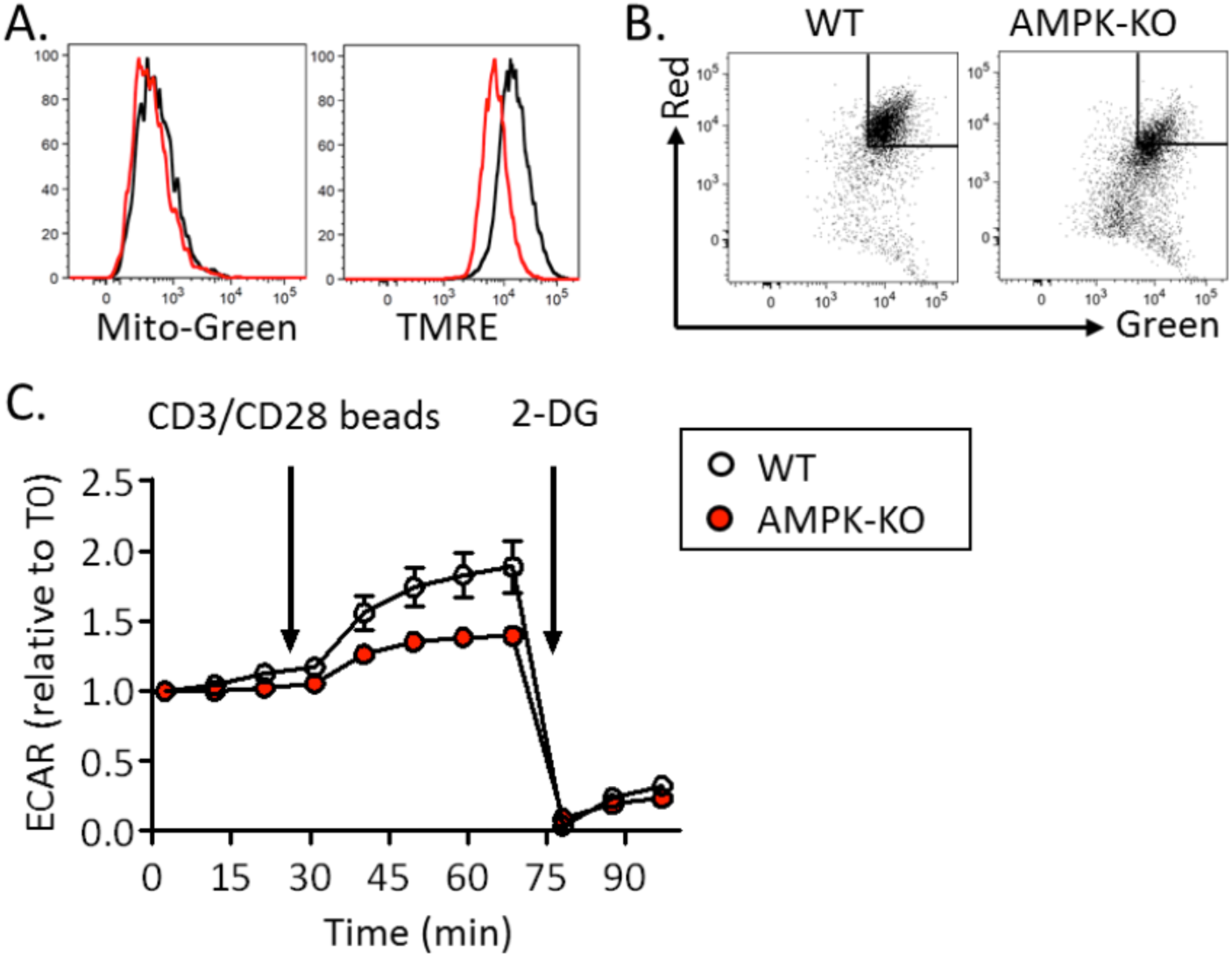
Defective mitochondrial stress-induced glycolytic switch in AMPK-KO memory-like Th cells. Naive CD4^+^ T cells from WT or AMPK^KO-T^ mice were polarized to memory-like Th1 cells by a 3-day stimulation in Th1 condition followed by an additional 4-day culture in IL-7-supplemented media. **(A)** Mitochondrial mass (Mito-Green) and **(B)** mitochondrial membrane potential (TMRE) in WT and AMPK-KO memory-like Th1 cells. Data are representative of 3 independent experiments. **(C)** ECAR measurement after treatment of memory-like Th1 cells with anti-CD3/CD28-coated beads as determined by Seahorse experiment. Data are representative of 2 independent experiments with n = 3.

**Figure S11:**
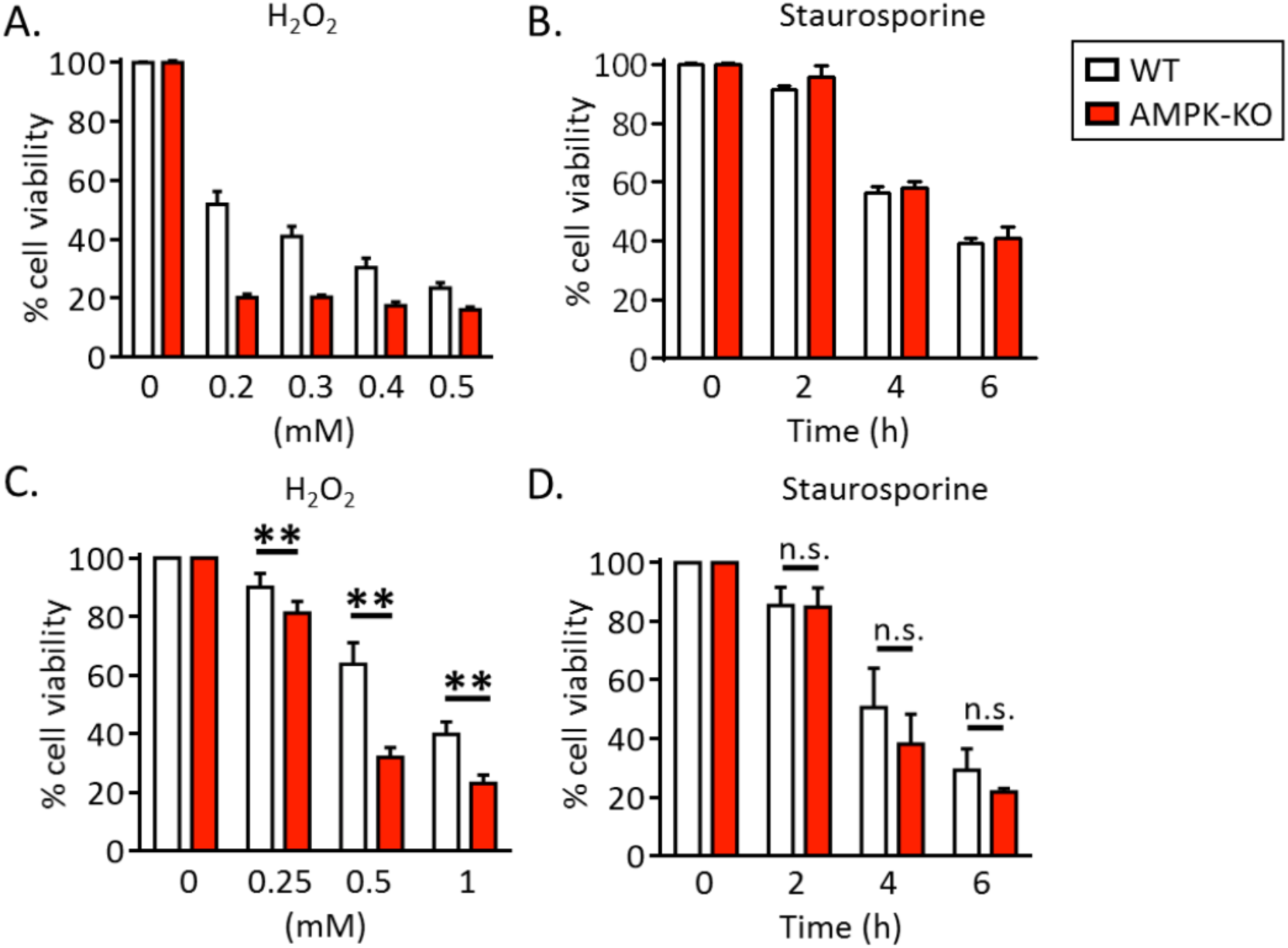
AMPK dampens ROS toxicity. Percentages of naïve T cell **(A, B)** or memory-like Th1 cell **(C, D)** viability after 2h-treatement with graded doses of H_2_O_2_ **(A, C)** or after treatment with staurosporine (4h, 5 µM) **(B, D).** Results are representative of 5 individual experiments (n = 2) or pooled from 4 (n = 4), 2 (n = 6) and 3 (n = 3) independent experiments **(B, C, D** panels, respectively). Statistical analysis: Mann-Whitney **(B-D).** ** p < 0,01

